# GBP1 drives lysosomal activity against cytosol-invading mycobacteria in macrophages

**DOI:** 10.1101/2025.04.28.650982

**Authors:** Mayra A. Aguirre-García, Sharon Portelli, Monica Varela, Annemarie H. Meijer

## Abstract

Guanylate-binding proteins (GBPs) are a family of interferon-inducible proteins playing diverse roles in the immunity to infections. By docking onto pathogens or pathogen-containing vesicles, GBPs can boost proinflammatory signaling. However, the interplay between GBPs and the phagolysosomal mechanisms mediating pathogen degradation remains largely unclear. Here, we elucidate the key role of GBP1 in driving lysosomal activity against mycobacterial infection in human macrophages. We show that phagosome damage induced by the mycobacterial ESX-1 secretion system triggers calcium release, and thereby the recruitment of GBP1, which resides in lysosomes upon interferon gamma activation. GBP1 targeting of mycobacteria-containing phagosomes leads to the recruitment of other GBPs but does not lead to the activation of membrane repair mechanisms or antimicrobial autophagy. Instead, GBP1 mediates lysosomal recruitment to intracellular mycobacteria, and promotes the maturation of phagolysosomes and their proteolytic activity required for the degradation of the mycobacterial cargo. Collectively, this work highlights the host-protective function of GBP1 in macrophages, where GBP1 acts as a key cellular sensor of mycobacterial infection and promotes the clearance activity of the host lysosomal system.

## INTRODUCTION

Autonomous immunity of macrophages plays a fundamental role in the host defense against mycobacteria, the causative agents of tuberculosis (TB) and other serious infections in humans and various animals species (Shamaei & Mirsaeidi, 2021; Smith, 2003) . During infection, macrophages are exposed to an environment of immunity-stimulating cytokines, among which interferon-gamma (IFNγ) is a key activator of microbicidal states (Ivashkiv, 2018). The IFNγ-inducible signals include a family of guanosine triphosphatases (GTPases), known as guanylate binding proteins (GBPs) (Tretina et al., 2019). Accumulating evidence in recent years has shown that GBPs marshal immunity to infection through several mechanisms, namely, by promoting interactions with the cellular machinery involved in cell death, by activating proinflammatory signaling pathways, and, in a less well understood manner, by triggering antimicrobial mechanisms, such as autophagy (MacMicking, 2012; Tretina et al., 2019). GBPs stand out as strongly induced factors in the IFNγ-mediated immune response to mycobacterial infection in macrophages (Rosain et al., 2023; Wei et al., 2022). Furthermore, in several clinical studies of TB and other mycobacterial diseases, GBPs emerge as a signature of the infection in terms of both transcriptional response and protein levels (Maertzdorf et al., 2011; Rosain et al., 2023; Teles et al., 2019; Yao et al., 2022). As a result, GBPs, notably GBP1, GBP4, and GBP5, have been proposed as new biomarkers for active TB (L. Chen et al., 2022; Shi et al., 2022; Yao et al., 2022). These clinical findings emphasize the need for mechanistic understanding of GBP function in the host defense to mycobacteria.

Following uptake by macrophages, pathogens are sequestered within phagosomes, which, in their mature state, fuse with lysosomes, a process eventually culminating in bacterial degradation (Gordon, 2016). However, many pathogens, notably pathogenic mycobacteria, have developed virulence mechanisms to subvert the phagolysosomal killing, either by inhibiting phagosome-lysosome fusion or by causing phagosome damage and gaining access to the cytosol (Chandra et al., 2022; Pauwels et al., 2017). As a cell-autonomous defense mechanism of macrophages, GBPs can bind directly on the surfaces of pathogens that have accessed the cytosol (Buijze et al., 2023; Feeley et al., 2017; Valeva et al., 2023). First, the binding of GBPs can disrupt the envelope function of gram-negative enterobacteria (Kutsch et al., 2020; Valeva et al., 2023; Zhu et al., 2024). Second, by interacting with lipopolysaccharide (LPS) on the surface of these bacteria, GBPs initiate a caspase-4-dependent inflammasome signaling platform resulting in cell death, thereby eliminating the pathogen’s intracellular niche (Feng et al., 2022; Fisch et al., 2020; Lagrange et al., 2018; Santos et al., 2020; Wandel et al., 2020). Other than directly interacting with cytosolic bacteria, GBPs also accumulate on pathogen-containing vacuoles, and this recruitment is dependent on the ability of pathogens to damage the phagosomal or phagolysosomal endomembranes (Bass et al., 2023; Feeley et al., 2017; Zwack et al., 2017). Although GBP recruitments have been reported for both human and mouse macrophages infected with *Mycobacterium tuberculosis* (Mtb) and *Mycobacterium bovis* (*M. bovis*), the mechanism by which GBPs detect the mycobacteria-containing vacuoles (MCV) and function in the cell-autonomous defense against these pathogens is incompletely understood (Buijze et al., 2023; Kim et al., 2011).

While triggering inflammasome activation is the best established interaction of GBPs with the host immune system, several studies have proposed GBPs to cooperate also with host proteins involved in autophagy and vesicular trafficking (Feeley et al., 2017; Haldar et al., 2014; Kim et al., 2011). Atg3 and Atg5 were found to be required for the proper binding of mouse Gbps to pathogen-containing vacuoles, suggesting that autophagy effectors might control GBP-dependent cell resistance to *Chlamydia trachomatis* and *Toxoplasma gondii* (Haldar et al., 2014). During Group A *Streptococcus* infection, human GBP1 contributes to a protective autophagy response, following membrane damage detection by galectin 3 (Gal3) (Hikichi et al., 2022). In addition, it was observed that mouse Gbp1 binds to the selective autophagy receptor Sqstm1/p62, resulting in the delivery of this complex to the *M. bovis*-containing phagosome (Kim et al., 2011). Further supporting the role of GBPs in host resistance, it has been reported that down-regulation of mouse Gbp2b resulted in a reduced autophagy response to *M. bovis* via the AMPK/mTOR/ULK1 pathway (Yu et al., 2022). Studies on *Leishmania* and *Chlamydia* infections suggest that mouse Gbps and human GBP1/2 promote fusion of autophagosomes and lysosomes with the pathogen-containing vacuoles (Al-Zeer et al., 2013; Haldar et al., 2020). However, the precise interaction of GBPs with the lysosomal system, responsible for bacterial degradation, is yet poorly understood.

Here, we studied the role of GBPs, especially GBP1, in the IFNγ-mediated immunity to mycobacteria in human macrophages. We used *Mycobacterium marinum* (Mm), a widely used surrogate model for Mtb (Guallar-Garrido & Soldati, 2024; Ramakrishnan, 2020; van der Vaart et al., 2014), to demonstrate that GBP1 is required for the innate immune response to this pathogen and to elucidate the underlying mechanism. We assessed the impact of the membrane-damaging ESX1 secretion system of Mm, showing that it is essential for triggering a robust calcium signaling response and for recruiting GBP1, which forms a complex with GBP2 and GBP4 on the MCV. While neither membrane repair nor autophagy responses were affected by loss of GBP1, we uncovered a GBP1-dependent mechanism whereby lysosomes are recruited to damaged phagosomes and the maturation and proteolytic activity of phagolysosomes is augmented by GBP1. Collectively, our results expand on the multifaceted ability of GBPs to regulate responses to intracellular pathogens, showing that GBP1 is involved in macrophage autonomous immunity by driving the lysosome system to damaged phagosomes and promoting crucial mycobactericidal functions.

## RESULTS

### The ESX-1 secretion system of Mm exhibits conventional virulence in human macrophages

Despite differences in transmission and tropism, Mtb and Mm share critical mechanisms in the pathogenesis within the host, which have led to the wide use of Mm as a surrogate model for Mtb (Stinear et al., 2008; Tobin & Ramakrishnan, 2008). The ESX-1 Type 7 secretion system is encoded by a genomic locus called ‘region of difference 1’ (RD1) and promotes phagosome membrane damage and access to the cytosol, a key feature of Mtb and Mm intracellular virulence (Carlsson et al., 2010; Hsu et al., 2003; Kurenuma et al., 2009; Lienard et al., 2020; Volkman et al., 2004) (Fig 1A). We therefore investigated whether Mm ESX-1 recapitulates the escape from phagosomes in INFγ-primed U937 macrophages and human monocyte-derived macrophages (hMDM). First, we assessed the kinetics of phagosome permeabilization using ubiquitin coating as marker for contact of bacteria with the cytosol upon infection (Campos et al., 2022), resulting in complete or partial coating of the bacterial surface (Fig 1A, Fig S1A). We observed that the ubiquitination level did not change significantly during the first 10 hours post infection (hpi). However, a slight peak was observed at 6 hpi (Fig S1B), and this time point was therefore selected to determine the role of ESX-1 in Mm accessing the cytosolic compartment. As expected, we found that 20 to 30 percent of wild-type (WT) Mm was ubiquitinated in both hMDM or U937 macrophages and that ubiquination was abolished in macrophages infected with the ESX-1-deficient Mm ΔRD1 mutant (Fig 1B and C, Fig S1 C). We then studied Gal3 and galectin-8 (Gal8), which are damage markers, known to be recruited after the exposure of glycosylated proteins by ruptured vesicles(Johannes et al., 2018) (Fig 1A). Similarly, we observed almost no colocalization of Gal3 or Gal8 in cells infected with the ΔRD1 mutant, while these markers colocalized with approximately 10 to 40 percent of WT Mm bacteria (Fig 1D-G, Fig S1 D and E). These results are consistent with what has been observed in other *in vitro* infection models of Mtb and Mm, and confirm the requirement of ESX-1 for cytosolic invasion of Mm in the U937 and hMDM cell types used in this study (Anand et al., 2023; Bernard et al., 2020; Osman et al., 2020; Stamm et al., 2003).

**Fig 1.**
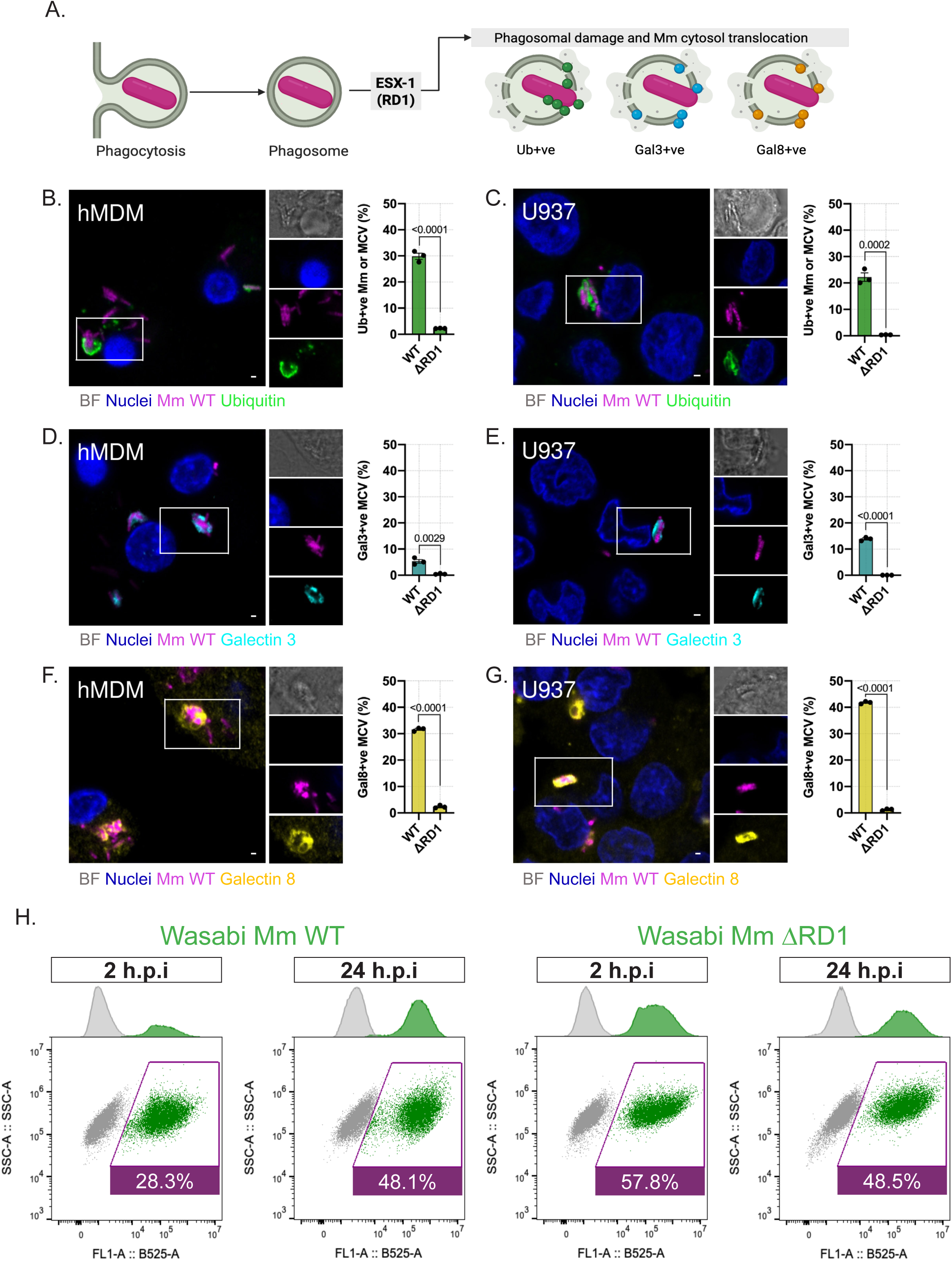
The ESX-1 secretion system of Mm is required for phagosome damage in hMDM and U937 human macrophages. **A.** Schematic illustrating ubiquitin (Ub+ve) and galectin (Gal3/8+ve) protein makers observed after rupture of a phagosome induced by the ESX-1 (RD1) secretion system in mycobacterial infections. **B-G.** ESX-1 requirement for ubiquitin, Gal3 and Gal8 recruitment to Mm infection. Representative single-stack confocal images (left) and quantification (right) from infected hMDM (B,D and F) and U937 (C,E and G) stained for ubiquitin (B and C), Gal3 (D and E) or Gal8 (F and G). Gray, Bright-field; blue, Nuclei; magenta, Mm Wild-Type (WT); green, ubiquitin; cyan, Gal3; yellow, Gal8. **H.** Representative fluorescence-activated cell sorting (FACS) dot plots of Mm Wild-Type (WT) and Mm ΔRD1 mutant (ΔRD1) at 2 h.p.i. and 24 h.p.i. Uninfected cells are in gray; infected cells are green. The manual gate strategy (magenta outline) and percentages of Mm positive cells are indicated in the dot graphs. Corresponding histograms are shown on top. *Data information*: graphs show percentages of ubiquitin (B and C), Gal3 (D and E) and Gal8 (F and G) positive Mm WT, Mm ΔRD1 or MCVs at 6 h.p.i, plotted as mean SEM from *n=*3 independent experiments with ≥ 200 cells in total per replicate and condition. p -values for indicated comparisons are shown on the graphs and calculated with t-test. Scale bars, 1µm.

We next sought to investigate the ability of U937 macrophages to phagocytose and restrict Mm WT and Mm ΔRD1 infection. At 2 hpi, we found a substantially higher number of cells infected with Mm ΔRD1 versus the WT strain (Fig 1H), suggesting a possible inhibitory effect of ESX-1 activity on phagocytosis. At 24 hpi, we observed that infection with the WT strain led to a 2-fold increase in bacterial load, whereas infection with the ΔRD1 strain had not increased since 2hpi, indicating that the presence of ESX-1 lowers the resistance capacity of the U937 macrophages (Fig 1H). These results support that the intracellular behavior of Mm in human macrophages matches key features of Mtb infection, showing that, Mm relies on ESX-1-dependent virulence for cytosolic invasion and intracellular growth (Bernard et al., 2020; Osman et al., 2020).

### GBP1 leads GBP2 and GBP4 recruitment to damaged Mm-containing vacuoles

GBPs are a family of INFγ-inducible proteins that have been implicated in cell-autonomous host defense to a large number of infections (Tretina et al., 2019). GBPs are specially known to target bacterial surfaces or pathogen-containing vesicles and induce cell death once the infection reaches the cytosol (Kirkby et al., 2023). We therefore hypothesized that GBPs would be recruited to the Mm bacteria or MCV. To test this, we performed immunohistochemistry on INFγ-primed hMDM and U937 macrophages infected with Mm WT at 6 hpi. We narrowed the analysis to GBP1, GBP2 and GBP4, for which antibodies are commercially available and previously validated (Bass et al., 2023; Valeva et al., 2023; Wandel et al., 2017). We observed that these GBPs were recruited to 10-25% of the Mm WT in hMDM or U937 macrophages (Fig2 A-F). In contrast, the recruitment of all three GBPs was almost abolished in hMDM or U937 macrophages infected with Mm ΔRD1 (Fig 2A,D,E, Fig S2 A-C). We therefore concluded that GBP1, GBP2 and GBP4 recruitment is dependent on the ESX-1 secretion system.

**Fig 2.**
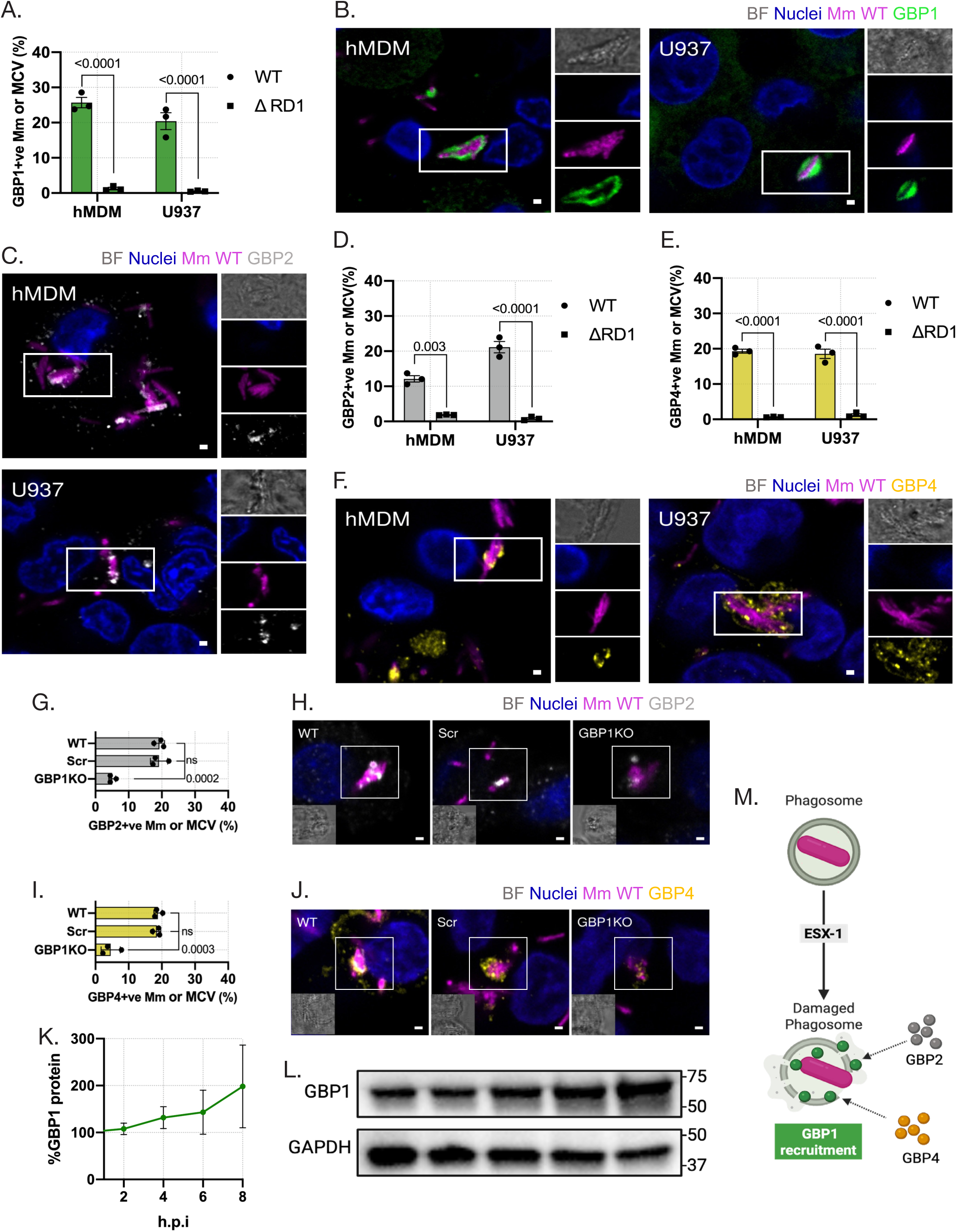
Mycobacteria recruit GBP1, GBP2 and GBP4 in an ESX-1 dependent manner. **A-F.** GBP1, GBP2 and GBP4 recruitment to WT and ΔRD1 Mm infection. Quantification (A,D and E) and representative single-stack confocal images (B,C and F) from infected hMDM (right) and U937 (left) stained for GBP1 (B), GBP2 (C) or GBP4 (F). Bright-field; blue, Nuclei; magenta, Mm Wild-Type (WT); green, GBP1; white, GBP2; yellow, GBP4. **G.H.** GBP2 positive Mm/MCV in the absence of GBP1. Quantification (G) and representative single-stack confocal images (H) from WT, Scr and GBP1KO U937 macrophages infected with Mm Wild-Type (WT) stained for GBP2. Bright-field; blue, Nuclei; magenta, Mm Wild-Type (WT); gray, GBP2. **I.J.** GBP4 positive Mm/MCV in the absence of GBP1. Quantification (G) and representative single-stack confocal images (H) from WT, Scr and GBP1^KO^ U937 macrophages infected with Mm Wild-Type (WT) stained for GBP4. Bright-field; blue, Nuclei; magenta, Mm Wild-Type (WT); yellow, GBP4. **K.L**. Western blot analysis of GBP1 protein levels during Mm infection. Quantification (right) and immunoblots (left) of GBP1 protein levels in WT, Scr and GBP1^KO^ U937 macrophages infected with Mm Wild-Type (WT). **M.** Summarizing scheme of dependency of GBP2 and GBP4 recruitment on ESX-1 and GBP1. *Data information*: graphs show percentages of GBP1 (A), GBP2 (D and G) and GBP4 (E and I) positive Mm/MCV of WT (A-J) or ΔRD1 infection(A, D and E) at 6 h.p.i, plotted as mean SEM from *n=*3 independent experiments with ≥ 200 cells in total per replicate and condition. p-values are indicated in the graphs and were calculated with Two-way ANOVA with Šidák multiple comparisons test (A,D and E) and One-way ANOVA with Dunnet’s multiple comparisons test (G and I). Scale bars, 1µm.

GBP1 has been demonstrated to lead the recruitment of other GBPs during bacterial infections such as, Shigella *flexneri* (S*. flexneri*), Salmonella *enterica* serovar Typhimurium (S. Typhimurium), and *Francisella tularensis* (F. *tularensis*) (Piro et al., 2017; Valeva et al., 2023; Wandel et al., 2017, 2020). Thus, we asked whether GBP1 is required upstream for the recruitment of GBP2 and GBP4 during Mm infection. To test this, we first investigated the co-recruitment of GBP1 with either GBP2 or GBP4. Among all GBP1 positive Mm in hMDM or U937 macrophages, only a subset was double positive for GBP2 or GBP4 (Fig S2 D-I), consistent with the idea that GBP1 recruitment is the upstream event. In agreement, we found that GBP2 positive Mm almost exclusively showed colocalization with GBP1, supporting that GBP2 recruitment relies on GBP1 (Fig S2 D-F). On the other hand, not all GBP4 positive Mm colocalized with GBP1 (FigS2 G-I). Next, we compared the recruitment of GBP2 and GBP4 in wild-type (WT), scramble (Scr) and GBP1 knock-out (GBP1^KO^) U937 macrophages. The Scr control contains a mutation in the OR5B17 gene, unrelated to infection, and serves to discriminate potential side effects from the CRISPR-Cas9 mutagenesis. We found that GBP2 and GBP4 recruitment in GBP1^KO^ U937 cells infected with Mm was strongly decreased but not entirely abolished (Fig 2G-J). Together, our results indicate that GBP1 promotes the recruitment of GBP2 and GBP4, but their colocalization with bacteria or damaged phagosomes does not exclusively depend on GBP1 during mycobacterial infection and might be tempo-spatially regulated.

Pathogens may impair the GBP-mediated cell-autonomous immunity by inducing the degradation of GBPs (Wandel et al., 2017). Therefore, we asked whether Mm infection might reduce GBP1 levels, and thereby inhibit its effector functions such as inducing GBP2 or GBP4 recruitment to Mm infection. We observed that protein levels of GBP1 in infected U937 macrophages did not decrease at different times post-infection implying that it is not depleted by the infection (Fig 2K and L).

Taken together, GBP2 and GBP4 recruitment during Mm infection is largely dependent on and preceded by GBP1 recruitment, and this process strictly depends on the Mm ESX-1 secretion system (Fig 2M).

### GBP1 colocalizes with ESX-1-damaged or sterile damaged endomembranes but does not promote galectin recruitment

GBP1 recruitment to pathogens exhibits notable plasticity. In some infections, GBP1 coats the pathogen-containing vacuole, while in others, GBP1 directly disrupts the bacterial membrane or binds to LPS, triggering inflammasome assembly. However, the activity against bacterial membranes has been investigated with gram-negative enterobacteria, which differ substantially in their physiology and pathogenicity from mycobacteria (Feng et al., 2022; Fisch et al., 2020; Kuhm et al., 2025; Santos et al., 2020; Valeva et al., 2023; Wandel et al., 2017). In fact, during Mm infection, we observed that human GBP1 displays recruitment patterns that suggest its association with endomembranes encapsulating one or more bacteria (Fig 3A). In agreement, we observed that at the majority of intracellular mycobacteria GBP1 overlaps with Gal8 signal marking endomembrane damage sites (Fig 3B and C). Based on these observations, we hypothesized that GBP1 targets damaged membranes of the MCV rather than cytosolic mycobacteria in macrophages. To validate this prediction, we treated macrophages and epithelial cells with l-leucyl-l-leucine methyl ester (LLOMe) to induce lysis of endolysosome membranes (Repnik et al., 2017). We found that GBP1-positive events were abundantly present after 30-60 min in hMDM macrophages, occurred to a lesser extent also in U937 macrophages, and were almost undetectable in the epithelial cells (HeLa) (Fig 3D and E). These results show that endogenous GBP1 responds to LLOMe treatment in macrophages and that it depends on the cell type or nature of the *in vitro* model how effectively GBP1 can sense intracellular membrane damage (Fig 3F).

**Fig 3.**
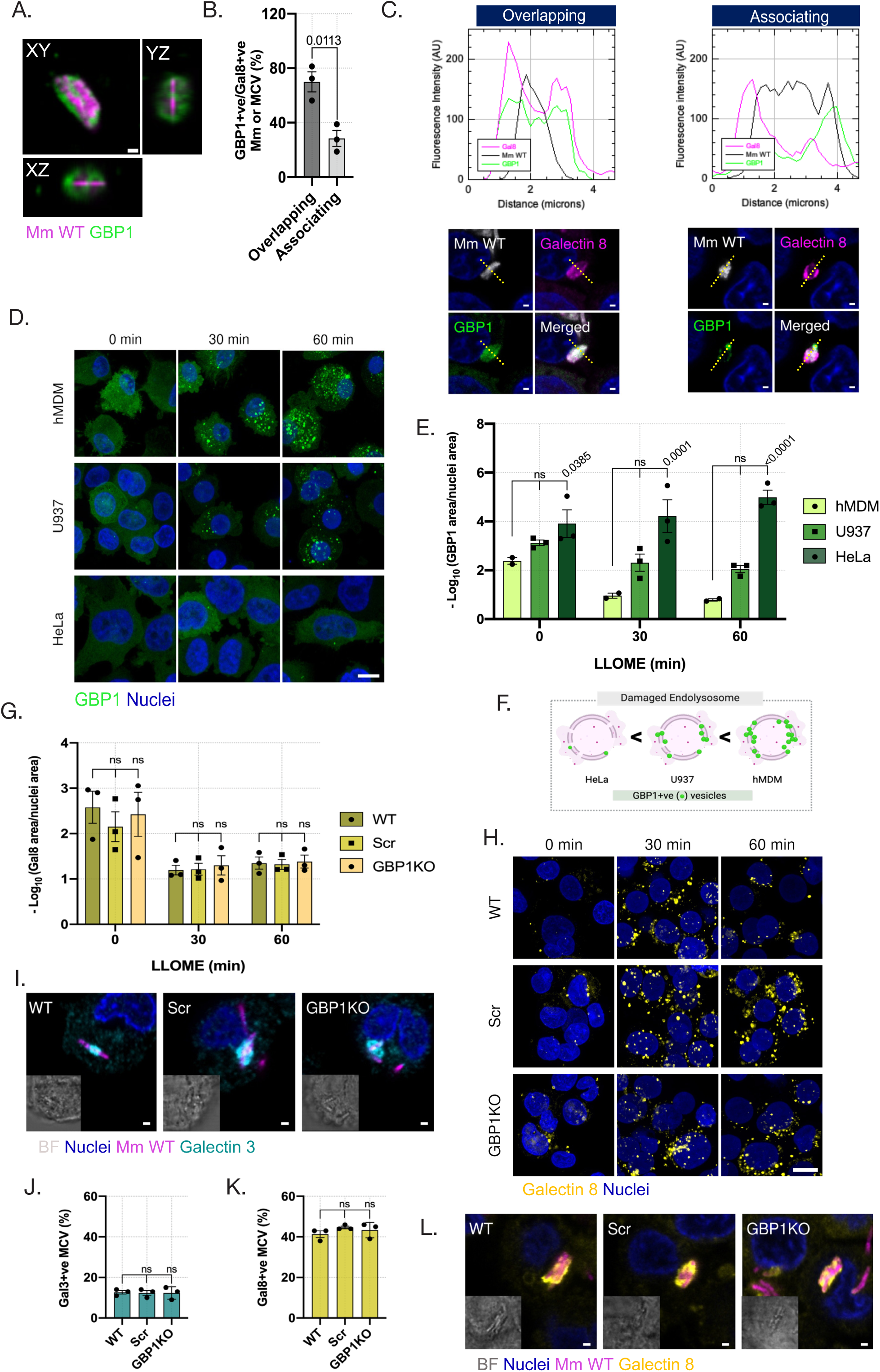
GBP1 colocalizes with damaged endolysosomes but does not promote Galectin recruitment. **A.** Phagosome-like GBP1 recruitment during Mm infection. Confocal image of GBP1 around Mm infection in hMDM. Orthogonal views are displayed to illustrate encapsulation of Mm by GBP1. Magenta, Mm Wild-Type (WT); green,GBP1. **B-C** GBP1 and galectin 8 co-recruitment to Mm infection. Quantification (B) of GBP1 and Gal8 positive MCVs when signals are overlapping or associating but not overlapping to the same MCV.Representative single-stack confocal images (C-bottom) from WT U937 macrophages infected with Mm Wild-Type (WT) stained for GBP1 and galectin 8. Gray, Mm Wild-Type (WT); Green, GBP1; Magenta, galectin 8. Profiles of fluorescence intensity (C-top) of Mm, GBP1 and Gal8 from the yellow transects represent the overlapping (left) and associating (right) patterns of both signals. **D-F.** GBP1 response to sterile endolysosome damage. Representative single-stack confocal images (D) from hMDM (top), U937 macrophages(middle) and HeLa cell (bottom) treated with LLOME and stained for GBP1 at 0 min, 30 min and 60 min post-treatment. Blue, Nuclei; green, GBP1. Quantification (E) of area corresponding to GBP1 positive vesicles normalized to nuclei area.Scheme (F) summarizing the potency of GBP1 recruitment to sterilely damaged endolysosomes among the three *in vitro* models. **G-H.** Gal8 response to sterile damage in GBP1-deficient macrophages. Quantification (G) of area corresponding to galectin 8 positive vesicles normalized to nuclei area. Representative single-stack confocal images (H) from WT, Scr and GBP1^KO^ U937 macrophages treated with LLOME and stained for Galectin 8 at 0 min, 30 min and 60 min post-treatment. Blue, Nuclei; yellow, Gal8. **I-L.** Gal3 and Gal8 responses to Mm infection in GBP1-deficient macrophages. Quantification of percentage of Gal3 (J) or Gal8 (K) positive WT Mm/MCVs from WT, Scr and GBP1^KO^ U937 macrophages at 6 h p.i. Representative single-stack confocal images from WT, Scr and GBP1^KO^ U937 macrophages infected with Mm Wild-Type (WT) and stained for Gal3 (I) or Gal8 (L). Gray, Bright-field; blue, Nuclei; magenta, Mm Wild-Type (WT); cyan, Gal3; yellow, Gal8. *Data information*: Area of positive GBP1 (E) and Gal8 (G) sterile-damaged endolysosomes was normalized to the nuclei area and plotted as the mean SEM of the negative logarithm. Percentages of galectin 3 (J) and galectin 8 (K) positive WT Mm/MCV at 6 h.p.i, were plotted as mean SEM from *n=*3 independent experiments with ≥ 200 cells in total per replicate and condition. p-values for indicated comparisons were shown on the graphs and calculated with t-test (C), Two-way ANOVA with Šidák multiple comparisons test (E and G) and One-way ANOVA with Dunnett’s multiple comparisons test (J and K). Scale bars, 10µm (D and H) and 1µm (A and B, I and L).

While GBP recruitment to damaged membranes is known to be reduced in galectin-deficient cells (Feeley et al., 2017), we considered that, vice versa, loss of GBP1 might affect the extend of membrane damage and ensuing galectin recruitment. However, LLOME induced similar levels of Gal8 recruitment at 30-60 min in WT, Scr or GBP1^KO^ U937 cells, indicating that GBP1 is not required for this response (Fig 3G and H). In addition, during Mm infection, no difference was found in either Gal3 or Gal8 recruitment between WT, Scr and GBP1^KO^ U937 macrophages at 6 h.p.i. (Fig 3I-L). Taken together, GBP1 is recruited to damaged endomembranes in macrophages and does not facilitate the recruitment of galectins in response to sterile or infectious triggers.

### GBP1 recruitment to ESX1-damaged phagosomes requires calcium signaling

Bacterial pathogens can activate the release of calcium (Ca^2+^) from intracellular stores into the cytosol (King et al., 2020). It has been shown that ESX-1 modulates intracellular Ca^2+^ fluxes during mycobacterial infection in macrophages (D. Chen et al., 2025; Francis et al., 2014; Yang et al., 2014). This led us to hypothesize that phagosome damage induced by ESX-1 triggers GBP1 recruitment through Ca^2+^ signaling. We used an oregon green labeled Ca^2+^ chelator, (BAPTA-AM), to monitor macrophages infected with Mm WT and Mm ΔRD1 mutant for approximately 4 hours. Accumulation of cytosolic Ca^2+^ (Ca^2+^_cyt_) was observed in the cells infected with Mm WT and remarkedly less in those infected with Mm ΔRD1, confirming the requirement of the ESX-1 virulence system (Fig 4A and B). Interestingly, Ca^2+^_cyt_ displayed different distribution patterns. We observed transient homogenous cytosol dispersion and either steady or transient concentration around MCVs (Supplementary Videos 1-4). To test if GBP1 recruitment depends on increased Ca^2+^_cyt_ levels, we incubated Mm-infected U937 macrophages with different modulators of Ca^2+^ signaling. In the presence of the Ca^2+^ chelator, BAPTA-AM, we observed a rapid impairment of GBP1 recruitment (Fig 4C and D). This result is consistent with the hypothesis that fluctuations of Ca^2+^_cyt_ can regulate GBP1 recruitment during Mm infection. Transient increases in Ca^2+^_cyt_ have been associated with activation of Ca^2+^ channels during membrane trafficking processes (Bagur & Hajnóczky, 2017). The transient receptor potential mucolipin 1 (TRPML1) is the major Ca^2+^ channel in phagosomes and lysosomes. We hypothesized that artificial stimulation of TRPML1 can increase GBP1 localization to the MCV. When Mm infected U937 macrophages were treated with a TRPML1 selective agonist (MLSA1), GBP1 recruitment consistently increased over time (Fig 4C and D). Finally, we tested if a cytoplasmic effector of Ca^2+^ signaling would mediate GBP1 aggregation. Calmodulin is a major Ca^2+^ sensor in non-muscle tissue cells, and the Ca^2+^-calmodulin axis has been related to INFɣ signaling activation (Nair et al., 2002; Villalobo & Berchtold, 2020). We found that GBP1 localization to MCVs was significantly lower in U937 macrophages after 30 min of treatment with the calmodulin inhibitor, calmidazolium (Fig 4C and D). Collectively, these results support that intracellular Ca^2+^ flux in response to ESX-1 mediated phagosome damage promotes GBP1 recruitment during mycobacterial infection (Fig4 E).

**Fig 4.**
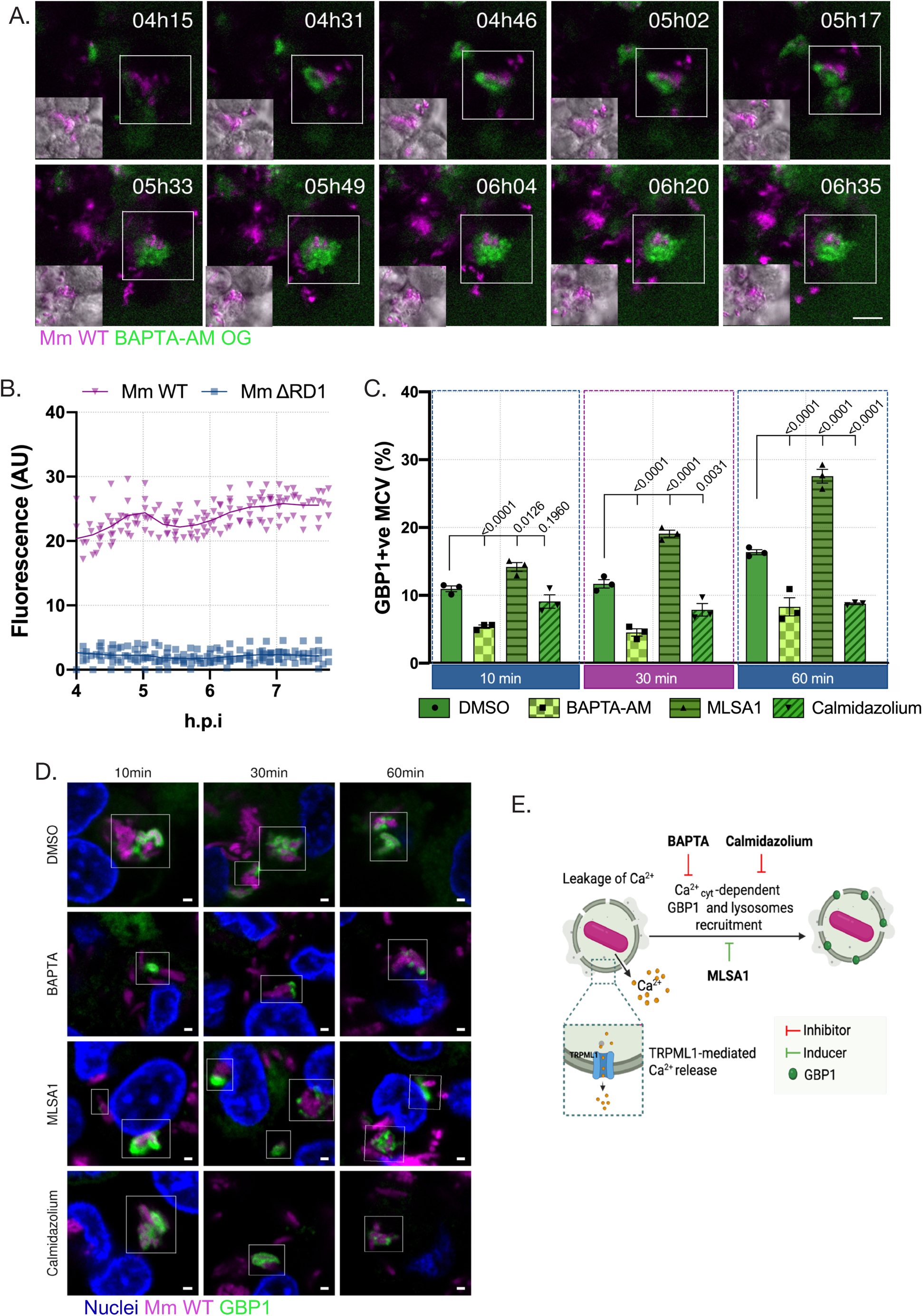
Mycobacterial damage of phagosomes increases cytosolic calcium and triggers GBP1 recruitment. **A-B.** Intracellular Ca^2+^ fluctuations in macrophages during mycobacterial infection. Representative single-stack confocal images from time-lapse microscopy (A) and quantification (B) of U937 macrophages infected with Mm Wild-Type (WT) and Mm ΔRD1 mutant (ΔRD1) stained with the membrane-permeable dye Oregon Green 488 BAPTA-AM (BAPTA-AM OG). Infected cells were imaged at 15-min intervals between approximately 4 and 8 h.p.i. Each dot (B) represents the BAPTA-AM OG fluorescence of an independent field (n=10) containing at least 20 cells normalized to the total cell area (µm^2^). Gray, Bright-field; magenta, Mm Wild-Type (WT); green, BAPTA-AM OG. **C-D.** Modulation of intracellular Ca^2+^ modifies the GBP1 recruitment rate to the MCV. Quantification (C) of GBP1 positive MCVs in U937 macrophages pre-infected for 4 h and treated with DMSO, BAPTA-AM (5µM), MLSA1 (20µM) and Calmodulin (10µM) for 10 min, 30 min and 60 min. Representative single-stack confocal images (D) from WT U937 macrophages infected with Mm Wild-Type (WT) and treated as above noted and stained for GBP1. Blue, Nuclei; magenta, Mm Wild-Type (WT); green, GBP1. **E.** Scheme summarizing the effects on GBP1 localization to MCVs during chemical manipulation of intracellular Ca^2+^. *Data information*: graph (A) shows fluorescence of BAPTA-AM OG is plotted as mean from n=10 independent fields with ≥ 200 cells in total. Percentages of GBP1 (C) positive MCVs at indicated time post-treatments were plotted as mean SEM from *n=*3 independent experiments with ≥ 200 cells in total per replicate and condition. p -values are indicated in the graphs and were calculated with Two-way ANOVA with Dunnett’s multiple comparisons test (C). Scale bars, 10 µm (A), 1 µm (D).

### Calcium-driven lysosomal fusion with Mm-containing phagosomes depends on GBP1

Ca^2+^ signalling has been associated with lysosome traffic towards the cargo-containing vesicles in a range of diseases, including infectious diseases (Li et al., 2016; Westman et al., 2020). Ca^2+^ signalling has also been implicated in the fusion of lysosomes with phagosomes-and autophagosomes-containing Mtb (Malik et al., 2000). However, the cellular events that might preceed or result from the vesicle trafficking are yet to be fully determined. We hypothesized that Ca^2+-^dependent recruitment of GBP1 to MCVs can facilitate lysosome fusion to restrict Mm growth and prevent invasion of the cytosol. We first asked whether Ca^2+^ signalling can regulate lysosome fusion during Mm infection, similar to GBP1 recruitment. To test this, we used lysosome-associated membrane protein 1 (LAMP1) to identify lysosomes, and analyzed its close association with MCVs (Fig S3). Inhibition of Ca^2+^ signaling with BAPTA-AM or calmodulin significantly impaired the recruitment of lysosomes to MCVs (Fig 5A and B). Conversely, LAMP1 association with MCVs was increased when we used MLSA1 to stimulate the TRPML1 Ca^2+^ channel, which is known as a key lysosomal fusion regulator (Ahuja et al., 2016) (Fig 5A and B). We next evaluated the role of GBP1 on the recruitment. As hypothesized, LAMP1 displayed a lower recruitment to MCVs when GBP1 was absent (Fig 5C and D). We considered that lysosome number can change to meet the demands of cells undergoing infection (Sachdeva & Sundaramurthy, 2020). At different times post-infection, we observed similar patterns of LAMP1 protein level between WT and Scr conditions, while GBP1^KO^ had a persistent lower LAMP1 level, reaching significance at 8 h.p.i (Fig 5E and F), suggesting that decreasing total levels of lysosomes might contribute to the effect of GBP1 on LAMP1 recruitment to MCVs. This result let us to assess the potential role of GBP1 in lysosomal biogenesis. We quantified the lysosome biogenesis master regulator, transcription factor EB (TFEB), whose activation induces its nuclear translocation. The nuclear compared to cytoplasmic levels of TFEB, however, appeared to be unrelated to GBP1 (Fig 5G and H). Taken together, these findings indicate that Ca^2+^ signalling regulates LAMP1 recruitment and the absence of GBP1 results in a decreased association of lysosomes with Mm-containing phagosomes.

**Fig 5.**
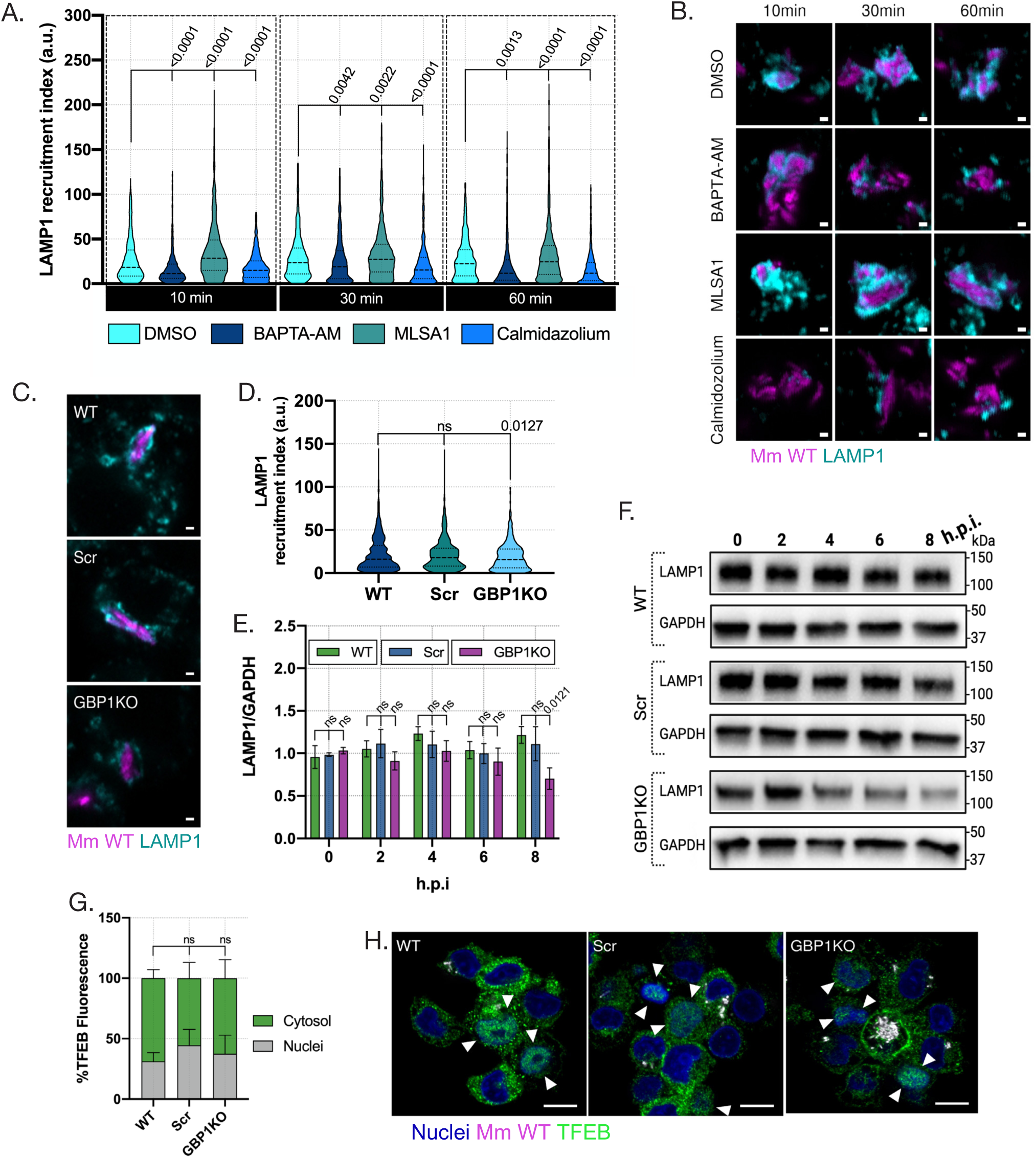
GBP1 promotes lysosomal fusion with Mm-damaged phagosomes. **A-B.** Modulation of intracellular Ca^2+^ modifies recruitment of lysosome marker LAMP1 to MCVs. Quantification (A) of recruitment index of LAMP1 to MCVs in U937 macrophages pre-infected for 4 h and treated with DMSO, BAPTA-AM (5µM), MLSA1 (20µM) and Calmodulin (10µM) for 10 min, 30 min and 60 min. Representative single-stack confocal images (B) from WT U937 macrophages infected with Mm Wild-Type (WT) and treated as above noted and stained for LAMP1. Magenta, Mm Wild-Type (WT); cyan, LAMP1. **C-D**. Lysosome recruitment to MCVs in the absence of GBP1. Quantification (D) of recruitment index of LAMP1 to MCVs from WT, Scr and GBP1^KO^ U937 macrophages at 6 h p.i. Representative single-stack confocal images from WT, Scr and GBP1^KO^ U937 macrophages infected with Mm Wild-Type (WT) and stained for LAMP1 (C). Magenta, Mm Wild-Type (WT); cyan, LAMP1. **E-F**. Analysis of LAMP1 protein levels in GBP1-deficient macrophages. Quantification (E) and immunoblots (F) of LAMP1 protein levels in WT, Scr and GBP1^KO^ U937 macrophages infected with Mm Wild-Type (WT) at different times post-infection. **G-H**. TFEB translocation to the nuclei during Mm infection in GBP1-deficient macrophages. Quantification of percentage of TFEB volume (G) in nuclei and cytosolic compartment in WT, Scr and GBP1KO U937 macrophages infected with Mm Wild-Type (WT). Representative single-stack confocal images (H) from WT, Scr and GBP1KO U937 macrophages infected with Mm WT stained for TFEB at 6 h.p.i. TFEB nuclei translocation showed in white squares (ROI). Blue, Nuclei; magenta, Mm WT; green, TFEB. *Data information*: graphs show recruitment index of LAMP1 (A) to Mm at indicated time post-treatments and in WT, Scr and GBP1KO U937 macrophages (D), plotted as mean SEM from *n=*3 independent experiments with ≥ 200 cells in total per replicate. Percentages of TFEB volume in nuclei and cytosol were plotter as mean SEM from *n=*3 independent experiments with ≥ 200 cells per replicate and condition. p-values are indicated in the graphs and were calculated with two-way ANOVA with Dunnett’s multiple comparisons test (E) and One-way ANOVA with Dunnett’s multiple comparisons test (A, D and G). Scale bar, 1µm (B and C) 10µm (H).

### GBP1 does not modulate the immediate autophagic response to Mm infection

Having shown that recruitment of lysosomes to intracellular Mm requires both Ca^2+^ signalling and GBP1 activity, we asked whether GBP1 would also be required for the anti-mycobacterial autophagy response. To this end, we first investigated the colocalization of GBP1 with ubiquitin, the initial signal to trigger antimicrobial autophagy. Ubiquitin signal intimately colocalized with GBP1 in hMDM and U937 macrophages infected with Mm (Fig 6A and B), suggesting that these proteins might share substrates and downstream roles. We thus examined whether this ubiquitin tagging was dependent on GBP1. No significant difference was observed between the controls and the GBP1^KO^ U937 macrophages regarding ubiquitination. Additionally, we analysed whether any other member of the GBP family was required for ubiquitin recruitment. Likewise, the levels of ubiquitinated Mm found in GBP2-5 ^KO^ cells did not show any significant difference when compared to those in WT macrophages (Fig6 C and D, S4). We next analyzed the formation of autophagosomes by immunostaining for LC3BI-II and found that GBP1^KO^ cells infected with Mm exhibited a similar percentage of Mm-LC3 positive autophagosomes as the WT and Scr controls (Fig 6E and F). Additionally, we quantified the levels of LC3B I-II displayed by WT, Scr, and GBP1^KO^ U937 macrophages over 8 h.p.i by western blotting. Mm-infected GBP1KO U937 cells showed a slight decrease, but not significant in the ratio of cytosolic and membrane-associated LC3B forms (Fig 6G and H). This result suggests that the autophagy flux is maintained. Collectively, GBP1 does not appear to modulate ubiquitination or autophagic targeting of Mm, suggesting that its antimicrobial function is mediated through lysosomes, independent of autophagy.

**Fig 6.**
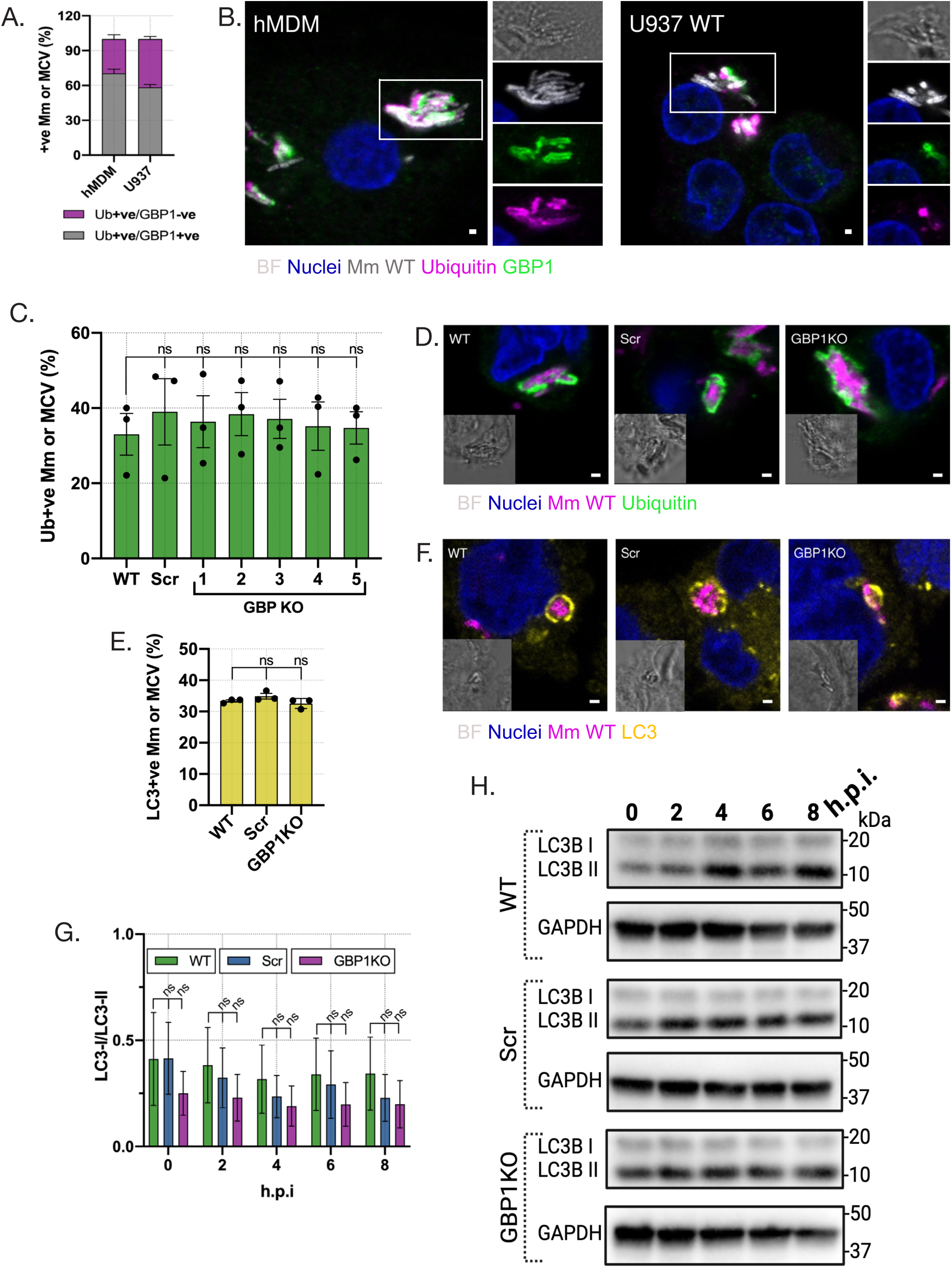
GBP1 does not modulate the immediate autophagic response to Mm infection. **A.B.** Colocalization of GBP1 with ubiquitinated Mm/MCV. Quantification (A) of percentage of GBP1 positive among ubiquitin positive Mm Wild-Type (WT) in infected hMDM and U937 macrophages at 6 h p.i. Representative single-stack confocal images (B) from hMDM (left) and U937 macrophages (right) infected with Mm WT and stained for GBP1. Gray, bright-field; blue, nuclei; white, Mm WT; ubiquitin; green, GBP1; magenta. **C-D.** Ubiquitination of Mm/MCV in GBPs-deficient macrophages. Quantification (C) of ubiquitin positive Mm Wild-Type (WT) Mm/ MCVs, and representative single-stack confocal images (D) from WT, Scr and GBP1-5^KO^ U937 macrophages stained for ubiquitin at 6 h.p.i. Gray, bright-field; blue, Nuclei; magenta, Mm WT; green, Ubiquitin. **E-F.** LC3-labelling of Mm or MCV in GBP1-deficient macrophages. Quantification (F) of LC3 positive Mm Wild-Type (WT) Mm/MCVs, and representative single-stack confocal images (G) from WT, Scr and GBP1^KO^ U937 macrophages stained for LC3 at 6 h.p.i. Gray, bright-field; blue, Nuclei; magenta, Mm WT; yellow, LC3B. **G-H**. Analysis of LC3 protein levels in U937 macrophages lacking GBP1. Immunoblots (H) and quantification (I) of LC3-II protein levels, relative to GAPDH, in WT, Scr and GBP1^KO^ U937 macrophages infected with Mm Wild-Type (WT) at different times post-infection. *Data information*: graphs show percentages of GBP1 positive ubiquinated WT Mm/ MCVs (A), ubiquitin (C) and LC3 (G) positive Mm WT at 6 h.p.i, plotted as mean SEM from *n=*3 independent experiments with ≥ 200 cells in total per replicate and condition. p -values are indicated in the graphs and were calculated with One-way ANOVA with Dunnett’s multiple comparisons test (A, C, G). Scale bar, 1µm.

### GBP1 resides in lysosomes upon INFγ induction

Based on our findings that lysosome recruitment requires GBP1, we next investigated to what extent lysosomes are associated with GBP1 positive MCVs. Our colocalization analysis showed that GBP1-LAMP1 closely localized during Mm infection (Fig 7A), raising the possibility that GBP1 is present on lysosomes before their fusion with MCV. The intracellular positioning of GBPs upon INFγ induction and before infection is not fully understood. We hypothesized that GBP1 resides in lysosomes after INFγ activation, which precedes the formation of GBP1-positive phagolysosomes after infection. To test this, we compared cellular fractions of untreated and INFγ-treated U937 WT macrophages under both uninfected and Mm-infected conditions. We analyzed four different fractions: Crude Cytoplasmic Fraction (CCF); Crude Lysosomal Fraction (CLF); Purified Lysosomal Fraction (PLF); and, Nuclei/Residual Fraction (NRF). As predicted, we found that endogenous GBP1 was only observed in INFγ-primed macrophage fractions. Confirming the presumed lysosomal localization, GBP1 was found in the CLF and PLF fractions together the lysosomal marker LAMP1. This lysosomal localization pattern of GBP1 was detected both in uninfected and infected conditions (Fig 7B).

**Fig 7.**
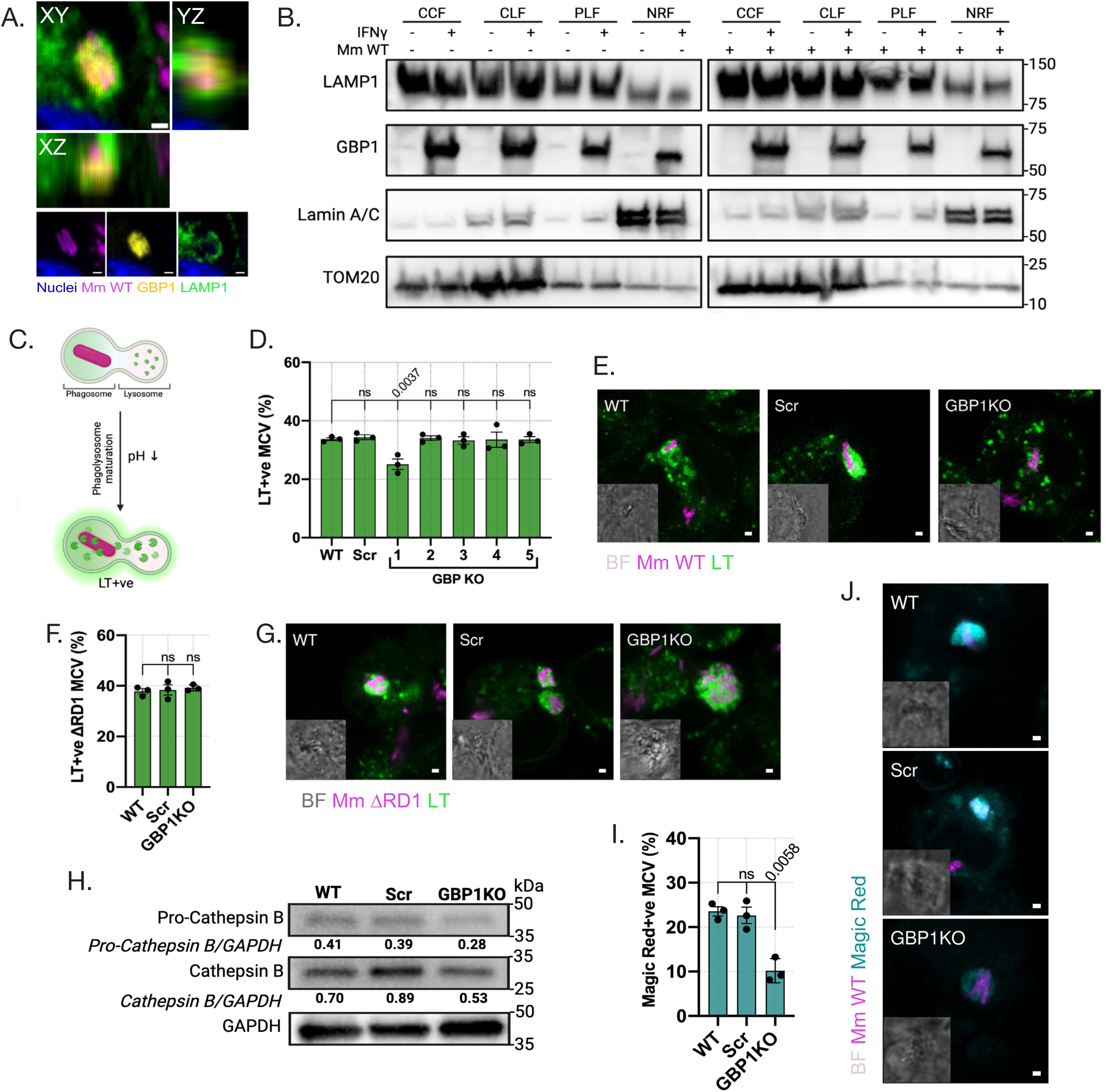
GBP1 resides in lysosomes and facilitates lysosome acidification and proteolytic activity. **A.** GBP1-LAMP1 colocalization with MCV. Orthogonal views are displayed to illustrate signal association of GBP1 and LAMP1 during Mm infection. Magenta, Mm Wild-Type (WT); yellow,GBP1; green,LAMP1. **B.** GBP1 presence in subcellular fractions rich in lysosomes in uninfected and infected conditions. Immunoblots showing GBP1 localization under the indicated conditions. Subcellular fractions: Crude Lysosomal Fraction (CLF); Purified Lysosomal Fraction (PLF); and Nuclei Residual Fraction (NRF) . Subcellular fractions were validated using different markers: LAMP1, lysosomal marker; LAMIN A/C, Nuclei marker; TOM20, mithocondrial marker. **C.** Scheme summarizing Lysotracker (LT) labelling of matured phagolysosomes undergoing a decrease of luminal pH. **D-E.** Acidification of mycobacteria in GBPs-deficient macrophages. Quantification (D) of percentage of Lyso-Tracker® (LT) positive MCVs, and representative single-stack confocal images (C) from WT, Scr and GBP1^KO^ U937 macrophages stained for LT at 6 h.p.i. Gray, bright-field; blue, Nuclei; magenta, Mm WT; green, LT. **F-G.** Acidification of Mm ΔRD1 mutant infection in the absence of GBP1. Quantification (E) of percentage of Lyso-Tracker® (LT) positive Mm ΔRD1 mutant (ΔRD1) MCV, and representative single-stack confocal images (F) from WT, Scr and GBP1^KO^ U937 macrophages stained for LT at 6 h.p.i. Gray, bright-field; blue, Nuclei; magenta, Mm ΔRD1; green, LT. **H**. Analysis of Cathepsin B protein levels in U937 macrophages lacking GBP1. Immunoblots and quantification of pro-Cathepsin and Cathepsin B protein levels in WT, Scr and GBP1^KO^ U937 macrophages infected with Mm Wild-Type (WT). **I-J.** Colocalization of cathepsin activity and mycobacteria in GBP1-deficient macrophages. Quantification (J) of percentage of Magic Red positive WT Mm/MCV, and representative single-stack confocal images (K) from WT, Scr and GBP1^KO^ U937 macrophages stained for Magic Red at 6 h.p.i. Gray, bright-field; blue, Nuclei; magenta, Mm WT; cyan, Magic Red. *Data information*: graphs show percentages of LT (D and G) and Magic Red (I) positive Mm WT or Mm ΔRD1 mutant at 6 h.p.i, plotted as mean SEM from *n=*3 independent experiments with ≥ 200 cells in total per replicate and condition. p -values are indicated in the graphs and were calculated with One-way ANOVA with Dunnett’s multiple comparisons test (D, G and H). Scale bar, 1µm.

### GBP1 regulates acidification and proteolytic activity of Mm-containing phagolysomes

Acidification is a critical process in lysosomal pathways, whereby the luminal pH is tightly regulated by the vacuolar ATPase (V-ATPase) to activate enzymatic functions of degradation. To investigate whether GBP1 can facilitate acidification of Mm in infected macrophages, we used LysoTracker (LT), which is able to diffuse into cells and fluorescently label acidic organelles such as lysosomes (Fig 7C). We determined the colocalization between Mm and acidic vesicles in WT, Scr and GBP1-5^KO^ U937 macrophages, and found that, out of all 5 GBP KO lines, only GBP1^KO^ U937 macrophages showed a significant reduction in the percentage of Mm colocalized with acidic vesicles (Fig 7D and E, S5 A). Next, we asked if ESX-1-dependent virulence is required for the effect of GBP1 deficiency on acidification. Indeed, LT signal was reduced in GBP1^KO^ U937 macrophages infected with WT Mm but not with the Mm ΔRD1 mutant (Fig 7 F and G).

Subsequently, we evaluated lysosomal protease activity driven by cathepsins (Yadati et al., 2020). First, we analyzed the protein level of Cathepsin B and found that GBP1 ^KO^ U937 macrophages displayed reduced total protein levels of pro-Cathepsin B and Cathepsin B when infected with Mm (Fig7 H). Using the Magic Red assay, we next observed that GBP1^KO^ U937 macrophages undergoing Mm infection displayed a reduced substrate cleavage, indicating lower proteolytic activity, which further confirmed that GBP1 mediates the functionality of lysosomes (Fig 7I and J). Finally, we analyzed total activity of Cathepsin B using live cell imaging. GBP1^KO^ U937 macrophages persistenly displayed lower Cathepsin B activity compared to WT and Scr conditions (Fig S5 B).

Collectively, these data support that GBP1 not only mediates the delivery of lysosomes to Mm in IFN-γ stimulated macrophages but also facilitates their degradative functions, including acidification and proteolysis.

## DISCUSSION

The lysosomal system is a necessary force in restricting intracellular infections by macrophages and is known to rapidly adapt its activity upon macrophage activation(Hipolito et al., 2018). Among the inducible signals downstream of macrophage activation by IFNγ, GBPs have emerged as a family of GTPases with diverse biological functions in the host immune response against intracellular bacteria (Kirkby et al., 2023; Meunier & Broz, 2016; Tretina et al., 2019). Clinical evidence has identified GBPs as markers of mycobacterial infections (L. Chen et al., 2022; Shi et al., 2022; Yao et al., 2022). Yet, if and how GBPs facilitate immunity against mycobacterial species remains largely unknown (Buijze et al., 2023; Kim et al., 2011; Olive et al., 2020). Here, we uncovered that, during mycobacterial infection, GBP1 is triggered by a Ca^2+^-dependent mechanism and functions as an activator of the lysosomal system downstream of the phagosomal damage caused by the pathogen, promoting the fusion of lysosomes with the MCV as well as subsequent acidification and proteolytic activity.

In bacterial infectious diseases, GBPs are mainly known to exert their functions when pathogens gain access to the cytosol . As shown here, Mm employs the ESX-1 secretion system, a virulence factor shared with Mtb, to damage the phagosome and subsequenly invade the cytosol of human macrophages. Using WT and ΔRD1 mutant bacteria, we show that ESX-1 is indispensable for GBP1, GBP2 and GBP4 recruitment during Mm infection. In agreement, Mtb-infected human macrophages displayed abundant GBP recruitment, while this response was absent during infection with *M. bovis* BCG, which is ESX1-deficient (Bass et al., 2023; Buijze et al., 2023; Feeley et al., 2017; Piro et al., 2017; Wandel et al., 2017). However, other studies showed that mouse GBPs can colocalize with M. bovis BCG and restrict its growth, which might suggest an ESX-1-independent damage response in mouse macrophages (Kim et al., 2011; Olive et al., 2020). Collectively, our results support that ESX-1 activity drives the hierarchical recruitment of GBPs during Mm infection, and, as in other bacterial infections, GBP1 functions as the main driver of this response (Britzen-Laurent et al., 2010; Santos et al., 2020; Wandel et al., 2017; Zhu et al., 2024). While ESX-1 activity elicits a GBP-mediated immune response, simultaneously, the ESX-1 virulence system could mediate the pathogenic evasion of GBP activity, as observed during Mtb infection in mice (Olive et al., 2020).

GBPs have been shown to bind directly to LPS on the surface of cytosol-invading gram-negative bacteria, triggering a signaling platform for caspase activation (Kuhm et al., 2025; Kutsch et al., 2020; Santos et al., 2020; Wandel et al., 2020; Zhu et al., 2024). However, there is currently no evidence that GBPs may bind directly to mycobacteria, which have a unique envelope in architecture and composition (Niederweis et al., 2010). In contrast, the ESX-1-dependent recruitment of GBPs and their colocalization with membrane damage markers as Gal3 and Gal8, favors that GBPs localize on damaged MCVs. This mirrors the situation in Legionella infection, where Gal3 and the Dot/Icm secretion system are required for GBP2 delivery to pathogen-containing vesicles (Feeley et al., 2017). Further supporting the recruitment of GBP1 to damaged membranes, we and others have shown that GBP1 can also accumulate on sterile-damaged endolysosomes in macrophages (Buijze et al., 2023; Feeley et al., 2017, this study). It is noteworthy that we did not observe GBP1 accumulation on damaged endomembranes in epithelial cells, suggesting that this function might be specific to the defense response of macrophages or other immune cells. Further research aimed to determine how different cell types instrumentalize the GBP sensing mechanisms will enhance the understanding of the cellular specificity in their effector functions and potential as antimicrobial agents.

How GBP1 activity is regulated by intracellular signalling is just beginning to be understood (Honkala et al., 2020; Kirkby et al., 2023). Under homeostatic conditions, membrane association of GBP1 is prevented due to its phosphorylation by the INFγ-induced kinase PIM1 (Fisch et al., 2023). Depletion of PIM1 during *T. gondii* infection liberates GBP1, enabling its recruitment to the parasitic vacuole. However, *S.* Typhimurium does not appear to follow the same mechanism, indicating pathogen-specificity in the regulation of GBP activity (Fisch et al., 2023). Here, we propose a model in which mycobacteria cause phagosomal damage, inducing the release of intraphagosomal Ca^2+^, which then triggers GBP1 recruitment. While the Ca^2+^ regulatory networks involved in the macrophagic response to endomembrane damage during infection are not yet completely elucidated, we and others find that bacterial arsenal such as the mycobacterial ESX-1 secretion system elevates cytosolic Ca^2+^ (Francis et al., 2014; this study). We now show that ESX-1 dependent Ca^2^ signaling is the mechanism underlying GBP recruitment to damaged MCVs. Mtb has been shown to disrupt not only the MCV but also to disturb ion permeability at the plasma membrane, demonstrated by Ca^2+^ influx, promoting cell death and release of proinflammatory cytokines (Beckwith et al., 2020; Toniolo et al., 2023; Yang et al., 2014). Our live cell imaging revealed both transient and persistent Ca^2+^ deposits in close vicinity of MCVs of WT but not ΔRD1 Mm, supporting that phagosome rupture releases intraluminal Ca^2+^ into the cytosol, similar to what has been shown in sterile-damaged endolysosomes (Duran et al., 2024; Herbst et al., 2020; Santos et al., 2020). Corroborating this finding, a very recent study reported that Ca^2+^ leakage from Mtb-containing phagosomes triggers LC3-lipidation, restricting further phagosome damage (D. Chen et al., 2025). Furthermore, a Ca^2+^-binding protein was found to associate with sterile-damaged- and Mtb-damaged vesicles in epithelial cells and human macrophages, respectively (Beckwith et al., 2020; Jia et al., 2020). The Ca^2+^-accumulation surrounding the MCV thus likely provides a signaling benchmark to induce GBP1 mobilization to the damaged endomembranes. Our results show that disruption or enhancement of Ca^2+^ signaling, using well-established chemical agents, modulated GBP1 recruitment in line with the proposed model. Further investigation of the molecular mechanisms underlying the Ca^2+^-GBP1 interaction will help to dissect the signaling networks controlling GBP1 function.

GBPs have been extensively studied in regulating canonical and noncanonical inflammasome activation (Feng et al., 2022; Fisch et al., 2020; Kim et al., 2016; Lagrange et al., 2018; Meunier et al., 2015; Santos et al., 2020; Wandel et al., 2020). In contrast, the efforts to elucidate their function in antimicrobial systems such as autophagy and lysosome-driven killing are still limited (Feeley et al., 2017; Haldar et al., 2014; Kim et al., 2011; Yu et al., 2022). Our findings indicate that Ca^2+^ signaling, elicited by phagosome rupture, promotes lysosome recruitment to Mm infection in a GBP1-dependent manner. In agreement, other studies have observed altered colocalization between pathogens and lysosomes under conditions of GBP1 deficiency or overexpression (Al-Zeer et al., 2013; Glitscher et al., 2021; Haldar et al., 2020). While it has been shown that mycobacterial lipids can inhibit phago-lysosomal fusion, the GBP1-mediated recruitment of lysosomes could function as an INFγ-stimulated mechanism to rewire the system and enhance the anti-mycobacterial response(Gutierrez et al., 2004). Transcriptional regulation by TFEB tightly couples activity of the lysosomal system with lysosome biogenesis (Settembre et al., 2011). Therefore, we cannot distinguish to what extent the effect of GBP1 may be attributed to reduced fusion of lysosomes with the MCV or reduced lysosome numbers and LAMP1 protein level in GBP1 deficient macrophages infected with Mm. However, the fact that GBP1 deficiency does not trigger TFEB activation, suggests that a possible effect on lysosomes biogenesis would be limited. Evident candidates to mediate the interaction of GBP1 with lysosomes would be the autophagy effectors, as interactions of ubiquitin-receptors and LC3 with GBP1 have been found in diverse infection models (Biering et al., 2017; Kim et al., 2011). However, our results show that GBP1 does not modify critical steps of the autophagic pathway such as ubiquitination, galectin recruitment or LC3 association (Campos et al., 2022). Thus, despite that a significant proportion of MCVs are LC3 positive, we found no evidence for an interaction between GBP1 and the autophagy machinery, and conclude that GBP1 acts at the level of lysosomal function during Mm infection.

Similar to Rab GTPases, which are involved in lysosomal maturation, GBP1 harbors a CaaX motif, which targets it for isoprenylation and membrane insertion (Modiano et al., 2005). Considering the effect of GBP1 on lysosomes, we investigated if GBP1 itself is present on these organelles. Indeed, we detected GBP1 on purified lysosome fraction after IFNɣ stimulation, suggesting that GBP1 could serve a direct bridging function for recruiting lysosomes to the MCV. This GBP1-mediated lysosome recruitment could serve different, not mutually exclusive, roles. Ca^2+^-mediated fusion of lysosomes has been shown to maintain the integrity of fungal-containing phagosomes (Westman et al., 2020), reminiscent of the well-studied patching function of lysosomes during plasma membrane repair (Andrews et al., 2014). Therefore, lysosome recruitment might contribute to repairing the damage inflicted to the MCV by ESX-1 activity. Although we do not exclude this possibility, we found no effect of GBP1 deficiency on the association of damage makers with the MCV. This led us to explore the impact of GBP1 on lysosome functionality. Our results revealed that GBP1 is required for lysosomal acidification and cathepsin activity, two crucial and synergizing functions of macrophages to control mycobacterial infection. Cathepsins are also targets of mycobacterial evasion mechanisms, which emphasizes the need for host mechanisms, like GBP1 activity, to ensure appropriate degradative capacity in the lysosomal system (Lu et al., 2025; Pires et al., 2016).

The function of GBP1 in driving cytosol-accessing mycobacteria to lysosomal degradation described in this study increases mechanistic understanding of the antimicrobial activity of the GBP family, which is associated clinically with mycobacterial infections. Strong induction of GBP1 and family members correlates with progression to the active stage of tuberculosis disease and could either represent a protective response or a pathological hallmark (L. Chen et al., 2022; Shi et al., 2022; Yao et al., 2022). We propose that activation of the lysosomal system by GBP1 serves to restrict intracellular growth during the early stages of mycobacterial disease. In contrast, at later stages of infection, the activation of inflammasome-mediated cell death by GBP1 might exacerbate infection, as mycobacteria are known to benefit from the induction of lytic host cell death programmes (Mahamed et al., 2017). By demonstrating the requirement of Ca²⁺ signaling for GBP1-mediated lysosome trafficking and functional activation, our study extends the immune repertoire of GBP1 and might drive new approaches in therapeutic development, which should aim to balance the different pro- and anti-mycobacterial roles of GBP1.

## MATERIALS AND METHODS

### Cell culture

U937 monocytes and hMDM were maintained in RPMI 1640 Medium -ATCC modification (ThermoFisher Scientific) supplemented with 10% v/v fetal calf serum (Sigma) at 37 °C in 5% CO2. U937 knock out cell lines for GBP1-5 and OR5B17 were previously reported (Valeva et al., 2023; Wandel et al., 2020). All cell lines used were tested negative for mycoplasma using Mycostrip ® (Invitrogen). Buffy coats were obtained from healthy anonymous donors (Dutch adults) after written informed consent (Sanquin Blood Bank, Amsterdam, The Netherlands). CD14^+^ monocytes in 50 ng/ml M-CSF (Proteintech) from whole blood of healthy donors were differentiated into macrophages for one week as previously described (Van Den Biggelaar et al., 2024). To induce macrophage differentiation, U937 monocytes were seeded 36h prior to infection in medium containing 100 ng/mL Phorbol 12-myristate 13-acetate (PMA). U937 macrophages or human primary macrophages were treated with human INFγ (R&D Systems) as previously described (Valeva et al., 2023), unless otherwise specified. For microscopic imaging experiments, cells were differentiated and infected in μ-Slide 8 Well slides (Ibidi).

### Bacterial infections

*Mycobacterium marinum* WT (Mstrain, ATCC BAA-535) and ΔRD1 strain expressing mWasabi (Takaki et al., 2013) were grown overnight at 28°C in Difco Middlebrook 7H9 broth supplemented with 10% albumin-dextrose-catalase (ADC) and 0.05% Tween 80 (Sigma-Aldrich) and hygromycin. The overnight culture was grown to an OD600 of 0.6 to 0.8. The bacteria were washed with phosphate-buffered saline (PBS) and resuspended in complete RPMI 1640 medium. The bacterial suspension was diluted to reach the desired multiplicity of infection (MOI), assuming that OD600=1 is equivalent to ∼10^8^ CFU/mL. For all U937 macrophage infections an MOI of 20 was employed. After Mm infection, the cells were incubated at 32°C. After 2 h, the cells were washed and replaced with complete RPMI medium supplemented with 30μg/ml gentamicin (Sigma-Aldrich) and incubated for 10 min to remove extracellular bacteria. Next, the medium was replaced with complete RPMI and cells were incubated at 32°C until the desired times post-infection.

### FACS

After infection, infected U937 macrophages were washed once with warm PBS, detached with trypsin-EDTA. Cells were next fixed in 4% paraformaldehyde for 30 min under rotation (100rpm). Cells were washed twice with PBS. All samples were analyzed on a CytoFLEX flow cytometer. Bacterial burden was determined with the signal intensity of the bacteria using the FlowJo software.

### Lysotracker staining

Acidic vesicles were stained by incubating cells in RPMI medium enriched with LysoTrackerTM Red DND-99 (Invitrogen) (1:2500) for 30min. Cells were washed twice with PBS prior to fixation with 4% paraformaldehyde in PBS. Cells were then washed three times with PBS.

### Immunofluorescence

At the desired time post-infection, cells were washed twice with PBS and fixed in 4% paraformaldehyde for 10 min. Cells were washed three times in PBS and then permeabilized and blocked with PBSB (PBS, 0.02% w/v saponin (Sigma-Aldrich), 2% w/v Bovine Serum Albumin (BSA) (Sigma-Aldrich)) for 30 min. Cells were then incubated for 2h at RT with the appropriate primary antibody (*see* supporting Table 1) diluted in PBSB. Cells were washed again three times with PBS and incubated for 2h at RT in the appropriate secondary antibody diluted in PBSB for 2h at RT. Cells were stained using NucBlue DAPI (invitrogen) when desired.

### Cathepsin B Assay (Magic Red)

Cathepsin activity was determined using the Magic^Red^ kit according to the manufacturer’s instructions (Abcam). Differentiated U937 macrophages were incubated in RPMI complete medium enriched with Magic Red substrate for 1h. Ten pictures were taken per condition within 1h post-fixation.

### Ca^2+^ signalling assays

U937 macrophages were cultured and infected as described above. Stock solutions of Ca**^2+^** signaling modulators were diluted in RPMI: BAPTA-AM (5μM) Tocris Cat No. 2787, MLSA1(20μM) Abcam Cat No. ab144622, Calmidazolium (10μM) TargetMol Cat No. T10667 and DMSO. Treatments were administrated to cells at 4 h.p.i. by replacing the medium and incubating as described above until the times specified. Similarly, Oregon Green 488 BAPTA-AM (5μM) MedChem Cat No. HY-104058 was used to measure the intracellular calcium with live-cell imagining.

### Microscopy

Cells were imaged using Leica confocal microscopy systems, TCS SP8 and Stellaris 5, using 40x oil-immersion objective (NA 1.30) and 20X (N.A 0.75), 60x and 40x oil-immersion objective (NA 1.40) respectively. Cells were imaged using Leica confocal microscopy systems, TCS SP8 and Stellaris 5, using 40x oil-immersion objective (NA 1.30) and 20X (N.A 0.75), 60x and 40x oil-immersion objective (NA 1.40) respectively.

For the time-lapse imaging, ten fields of view containing more than >300 cells were imaged at intervals depending on the experiment: *intracellular calcium assay* (approx. 15 min), *Magic Red assay* (approx. 17 min). Fields of view were spread all over the slide based on the coordinates of each condition and using only channels corresponding to bright-field and bacteria to avoid bias.

### Image analysis

Fiji software from the ImageJ package (Schneider et al., 2012) was used to analyze the confocal images. For image analysis, Mm observations were defined as one or more bacteria in direct contact with each other (regardless of localization in the cytosol or MCV). The Z-step size was set between 0.5μM and 1μM. To quantify colocalization of Mm or MCV with vital dyes or with antibody signals, positive Mm or MCVs were counted through manual analysis of z-stack confocal slices (without projection method) unless specified otherwise.

#### Sterile damage of endolysosomes

The images from GBP1 and Gal8 positive endomembrane after LLOME treatment were processed with subtract (value=50) and Gaussian Blur filters (0.5-μm radius). A threshold was manually set to segment the fluorescence of either GBP1 or Gal8. The z-stacks were then projected into one image using a maximum-intensity projection. Total area was measured using analyze particles and normalized with the nuclei are per image. Finally, the negative logarithm base 10 was calculated.

#### Intracellular Ca^2+^ during Mm infection (live imaging)

The images from Mm-infected macrophages treated with BAPTA-AM OG 488 were processed by segmenting the cell area using the brightfield (BF) channel. BF z-stacks were firstly projected into one image using Standard deviation projection. Next, the contrast was enhanced (0.5%) and Gaussian Blur filter (0.5-μm radius) was applied. The segmented cell area was then obtained with a pre-established threshold, which maintained in all images and used for particle analysis. The masking was next transferred to a maximum projection image of the BAPTA-AM OG 488 channel. Total integrated intensity of chelated *Ca^2+^* was normalized to the area of macrophages.

#### LAMP1 recruitment to MCVs

The images from LAMP1 immunostainings were processed with Gaussian Blur filter (0.5-μm radius) on each channel (Mm and LAMP1). Next, bacteria were segmented by using the same manually-preestablished threshold in all conditions and all particles were analyzed across the z-stacks. The ROIs obtained from analyzed particles were next combined to connect particles belonging to the same MCV (objects). The masking was then expanded to measure the lysosome recruitment around Mm. Finally, the masking was transferred to the LAMP1 channel determining integrated intensities and areas across the z-stacks. Finally, an index of LAMP1 recruitment was calculated by the total sum of all integrated intensity values divided by all integrated area values across multiple stacks. Each *I_i_* and *A_i_* corresponds to the integrated intensity and area respectively of the *i*-th stack. The index starts at 1 and increases up to *n*, where *n* is the total number of stacks (Fig S3)

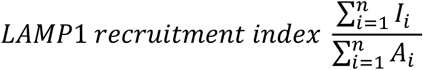

#### Magic Red (live imaging)

The activity of Cathepsin B was determined with Magic Red assay. Images from Mm-infected macrophages treated as described above, were processed by segmenting the bacteria and measuring the Magic Red fluorescence in the surrounding area. To exclude extracellular bacteria, the background from the cell channel (post-segmentation) was subtracted from the bacteria channel. The z-stacks of the bacteria channel were then projected into a single image using maximum projection. Then bacteria were segmented with a pre-established threshold, which was maintained in all images. The masking was next transferred to a maximum projection image of the Magic Red channel to determine fluorescence values and areas over time. The total integrated intensity of the Magic Red was normalized to the Mm expanded area.

### Lysosome isolation

Lysosomes from U937 macrophages were isolated using a lysosome enrichment kit according to the manufacturer’s manual (Sigma-Aldrich, LYSISO1). In brief, 1.5-2x10^8^ cells were differentiated and infected as described above. Cell were trypsinized and harvested. The packed cells were resuspended into extraction buffer and broken with a 7mL Dounce homogenizer using a Pestle B. 40 manual strokes were sufficient to achieve 95% of breakage (confirmed using Trypan blue). To obtain the Crude Cytoplasmic Fraction (CCF) (supernatant) and Nuclei Residual Fractions (NRF) (precipitate) samples were separated by centrifugation (1000 g x 1min). Next, to obtain the Crude Lysosomal Fraction (CLF) which still contains a mixture of light mitochondria, lysosomes, peroxisomes and rough endoplasmic reticulum (ER), the CCF was separated by centrifugation (22000 g x 20min). To obtain the Purified Lysosomal Fraction (PLF), the CLF was diluted to a solution that contains 19% Optiprep^TM^ density gradient medium, followed by CaCl_2_ (8μM) precipitation, thereby removing ER and mitochondria and enriching for lysosomes.

### Western Blot

After the specified infection or treatments, U937 cells were washed with PBS (Sigma) and lysed by agitation for 30 min on ice using cold RIPA Lysis and Extraction Buffer (Thermo ScientificTM) with cOmpleteTM Mini EDTA-free Protease Inhibitor Cocktail (Roche). The protein concentration of the cell lysates obtained was determined using BCA Protein Assay Kit (Thermo Scientific) and subsequently the lysates were diluted using MilliQ water to normalize protein concentrations across different samples for comparability. The samples were then loaded onto Mini-PROTEAN® TGXTM Precast Gels (Bio-Rad) and transferred to 0.2μm PVDF Transfer Membrane (Thermo ScientificTM) for immunoblotting. Membranes next were blocked in 5% skimmed milk prepared in Tris-buffered saline containing 0.1% Tween 20 for 1 h. Primary antibody (*see* supporting Table 1) incubations were performed overnight at 4 °C in blocking solution. Membranes were washed three times with TBS-T for 5min. Secondary antibodies (see supporting Table 1) were then incubated for 2h at RT in blocking solution. Digital images were acquired using the Bio-Rad Universal Hood II imaging system (720BR/01565 UAS). Band intensities were quantified by densitometric analysis using Image Lab Software (Bio-Rad, USA) and values were normalized to GADPH as a loading control.

### Statistical analysis

Detailed statistical analyses for individual experiments are given in each figure legend on the *Data information* section. This includes the statistical test and its post-hoc correction (if applicable), and the numbers of cells, replicates and independent experiments accordingly to the experimental design. Data are given as means ± SEM and analyzed using two-sided unpaired t-test (*two groups*) or one-way ANOVA and two-way ANOVA with multiple comparisons (*more groups*). GraphPad Prism 8.0 was used to perform all statistical analyses. Statistical significance was set at p<0.5. All p-values are shown on the graphs, otherwise ns: not significant.

## Data availability

The codes used for analyzing the experiments are available on a GitHub repository at https://github.com/aguirregarciama/GBP1_coordinates_Lysosome_activity. The repository of the confocal imaging is available at BioImage Archive: <u>10.6019/S-BIAD1876</u>.

## Acknowledgments

We would like to thank Prof Dr. Thomas Henry and Dr. Brice Lagrange (CIRI, CNRS UMR5308 - Inserm U1111 - ENS Lyon - UCBL) for sharing the GBP1-5 KO U937 cell lines. We also thank Dr. Anno Saris (Leiden University Medical Center) for providing primary human macrophages. We are grateful to Dr. Joost Wilemse and Dr. Gabriel Forn Cuní (Institute of Biology Leiden) advise on image analysis and to Maria Parlani and Prof. Dr. Peter Friedl (Radboud University Medical Center, Nijmegen) for facilitating FACS analysis. M.A.A-G was funded by INFLANET, European Union’s Horizon 2020 research and innovation programme under the Marie Skłodowska-Curie grant agreement No 955576. M.V. was funded by the Dutch Research Council (NWO) Talent Scheme, Veni fellowship VI.Veni.192.151.

## Author contributions

**M.A.A-G**: conceptualization; data curation; formal analysis; investigation; methodology, ; visualization; writing – original draft; writing – review and editing. **S.P**: investigation; formal analysis; writing – review and editing. **M.V.:** conceptualization; funding acquisition; methodology; supervision; writing – review and editing. **A.H.**: conceptualization; data curation, funding acquisition; supervision; writing – review and editing.

---

**Fig S1.**
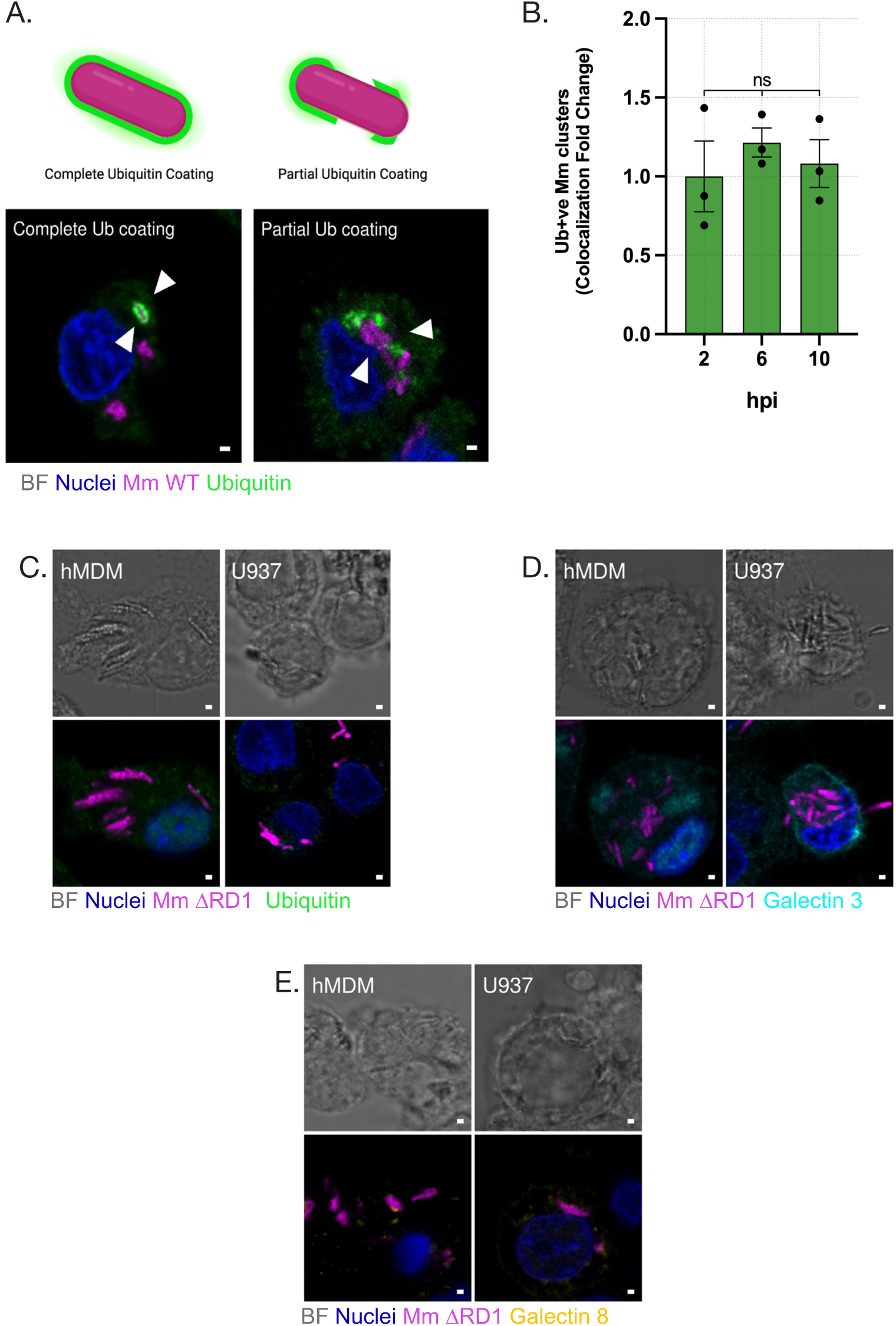
Supplementary data related to Fig 1. **A.** Schematic illustrating (top) and representative single-stack confocal images (bottom) of ubiquitin recruitment patterns to Mm infection. Gray, Bright-field; blue, Nuclei; magenta, Mm Wild-Type (WT); green, ubiquitin. Scale bar, 1µm. **B.** Quantification of fold change of ubiquitin recruitment to Mm WT or MCV at 2, 6 and 10 h p.i. Plotted as mean SEM n=3 independent experiments with ≥ 200 cells per replicate and condition. ns, not significant. p -value calculated with One-Way ANOVA with Tukey’s multiple comparisons test. **C-E.** Lack of recruitment of damaged-phagosome markers to Mm ΔRD1 mutant infection. Representative single-stack confocal images from infected hMDM (left) and U937 (right) stained for ubiquitin (C), Gal3 (D) and Gal8 (E). Gray, Bright-field; blue, Nuclei; magenta, Mm ΔRD1 mutant (ΔRD1); ubiquitin, green; Gal3, cyan; Gal8, yellow. Scale bar, 1µm.

**Fig S2.**
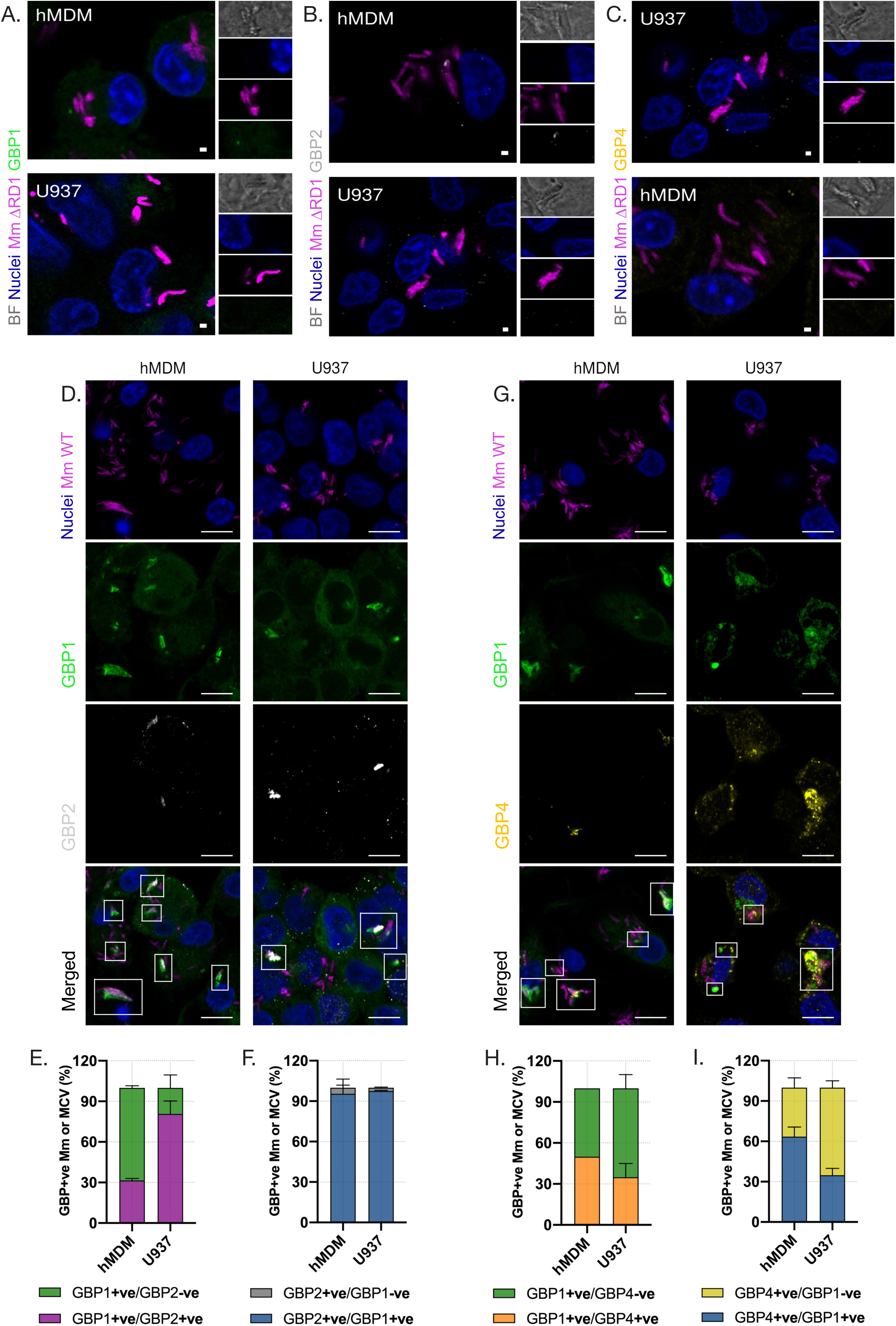
Supplementary data related to Fig 2. **A-C.** Limited recruitment of GBP1, GBP2 and GBP4 during Mm ΔRD1 mutant infection. Representative single-stack confocal images from infected hMDM (top) and U937 (bottom) stained for GBP1 (A), GBP2 (B) or GBP4 (C). Gray, Bright-field; blue, Nuclei; magenta, Mm ΔRD1 mutant (ΔRD1); GBP1, green; green, GBP1; white, GBP2; yellow, GBP4. Scale bar, 1µm. **D-F.** Colocalization of GBP1 with GBP2 in Mm-infected macrophages. Representative single-stack confocal images from infected hMDM (left) and U937 (right) stained for GBP1 and GBP2 (D). Examples of colocalization are outlined in the merged images. blue, Nuclei; magenta, Mm Wild-Type (WT); green, GBP1; white, GBP2. Quantification GBP2 co-recruitment with GBP1 during Mm infection. Percentage of GBP2-positive (E) Mm/MCV among GBP1-positive Mm/MCV and vice-versa (F). **G-I.** Colocalization of GBP1 with GBP4 in Mm-infected macrophages. Representative single-stack confocal images from infected hMDM (left) and U937 (right) stained for GBP1 and GBP4 (D). Examples of colocalization are outlined in the merged images. blue, Nuclei; magenta, Mm Wild-Type (WT); green, GBP1; yellow, GBP4. Quantification GBP2 co-recruitment with GBP1 during Mm infection. Percentage of GBP4-positive (H) Mm/MCV among GBP1-positive Mm/MCV and vice-versa (I). *Data information*: graphs show percentages of GBP1 (E-I), GBP2 (E and F) and GBP4 (H and I) positive Mm WT or MCV at 6 h.p.i, plotted as mean SEM from *n=*2-3 independent experiments with ≥ 200 cells per replicate and condition. Scale bar, 1µm (A-C) Scale bar, 10µm (D and G)

**Fig S3.**
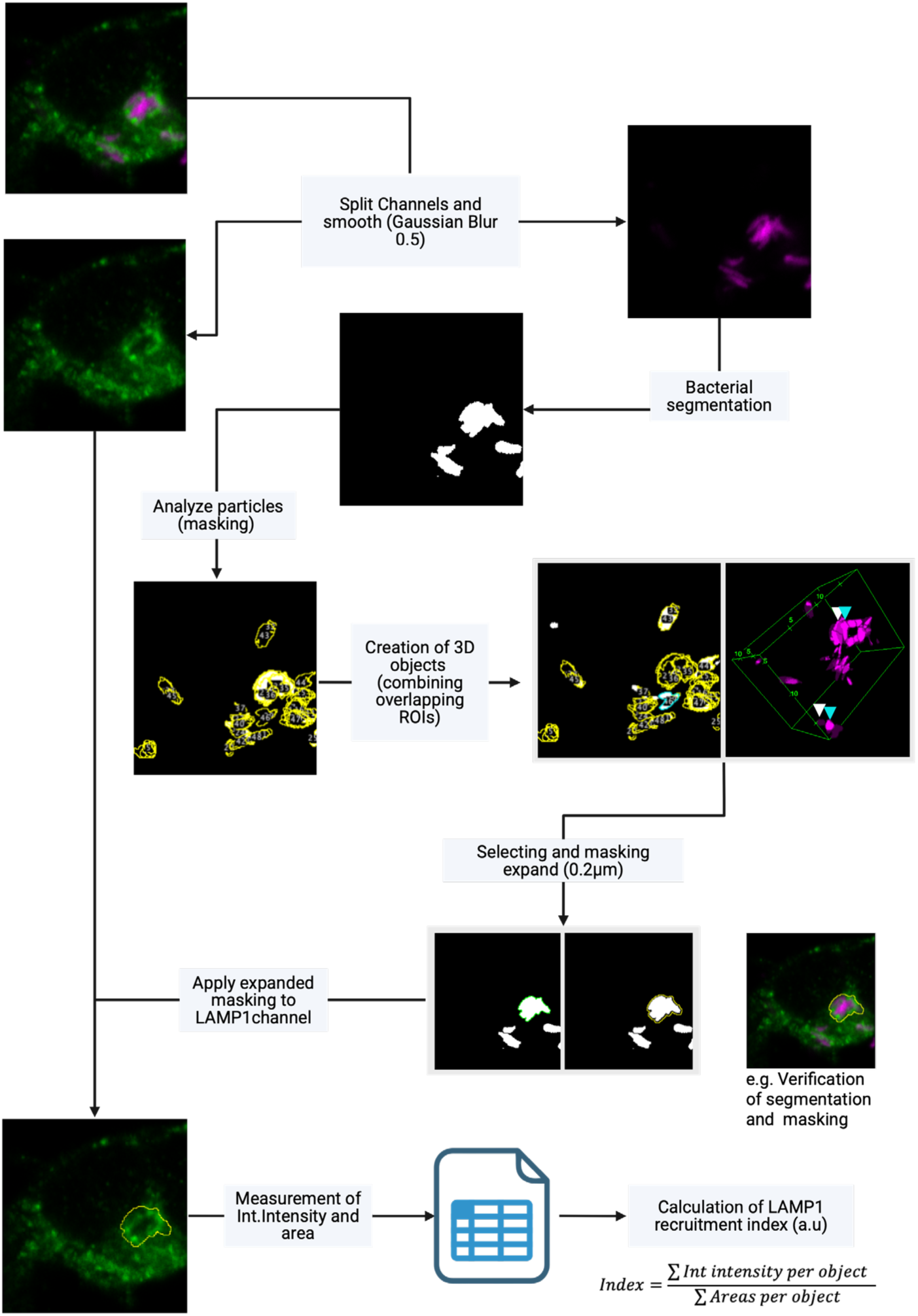
Supplementary data related to Fig 5. Diagram of image analysis workflow for the quantification of LAMP1 recruitment in response to mycobacterial infection in U937 macrophages.

**Fig S4.**
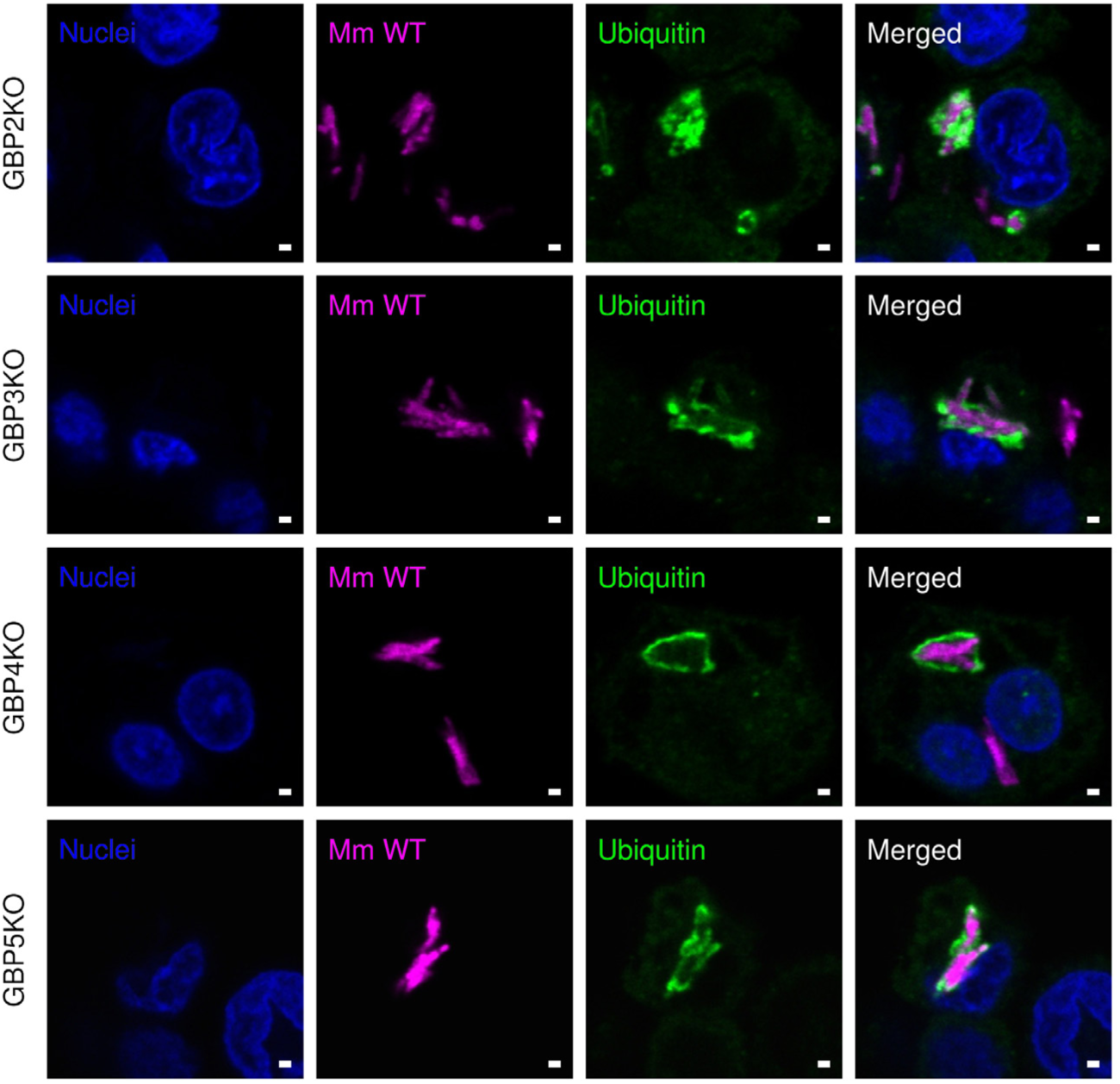
Supplementary data related to Fig 6. Ubiquitination of mycobacteria in GBP2-5^KO^ macrophages. Representative single-stack confocal images from Mm Wild-Type (Mm WT) infected GBP2-5^KO^ U937 macrophages and stained for ubiquitin. Gray, Bright-field; blue, Nuclei; Mm WT, magenta; ubiquitin, green. Scale bar, 1µm.

**Fig S5.**
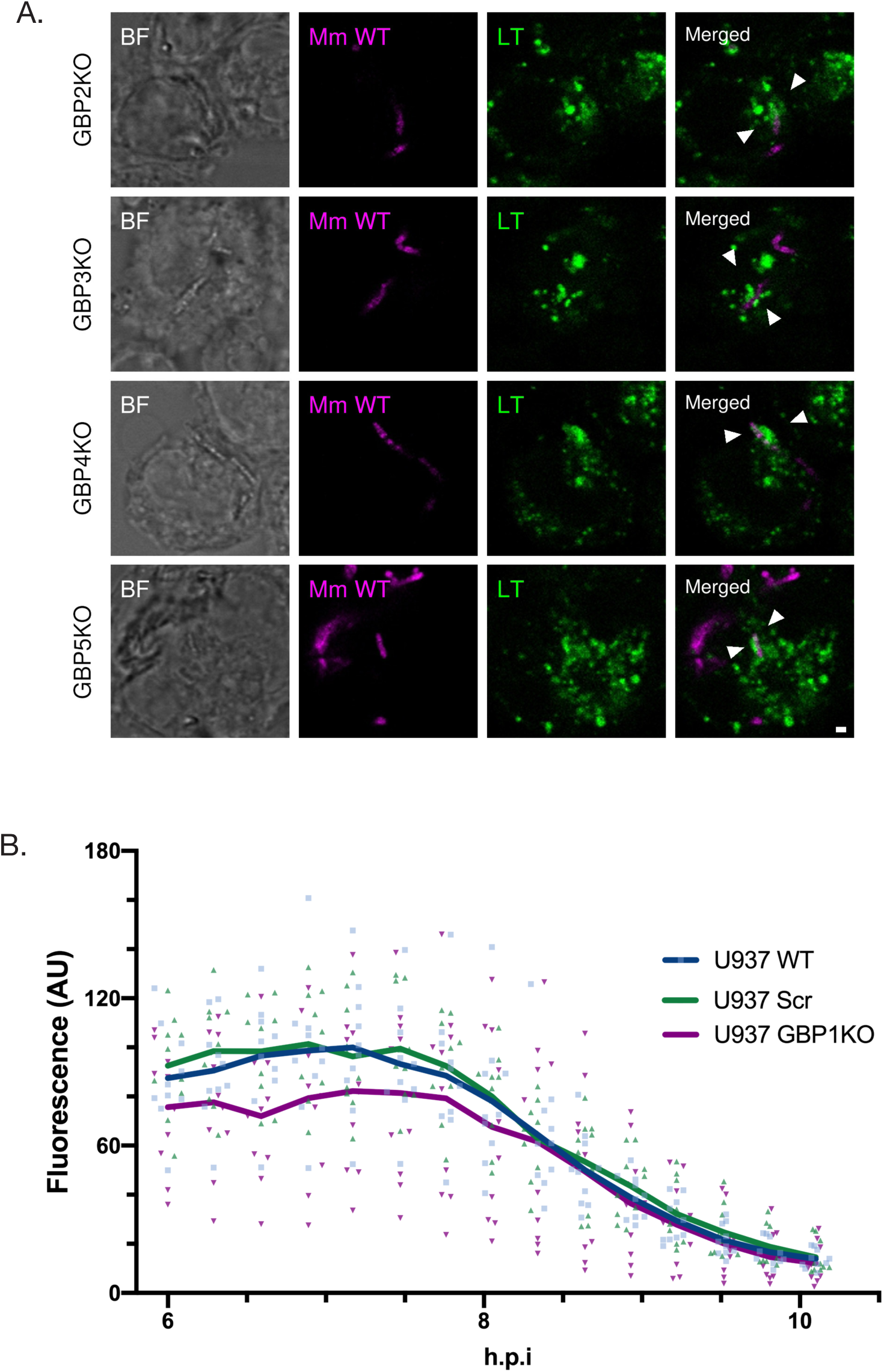
Supplementary data related to Fig 7. **A.** Acidification of mycobacteria in GBP2-5^KO^ macrophages. Representative single-stack confocal images from Mm Wild-Type (Mm WT) infected GBP2-5^KO^ U937 macrophages and stained for LysoTracker®(LT). Gray, Bright-field; blue, Nuclei; Mm WT, magenta;LT, green. Scale bar, 1µm. **B.** Activity of cathepsin B associated with mycobacteria in the absense of GBP1. Quantification of the Magic Red fluorescence in the Mm surroundings from WT, Scr and GBP1^KO^ U937 from 6 h.p.i.

**Supplementary video 1.** Transient accumulation of intracellular calcium around the MCV in U937 macrophages

**Supplementary video 2.** Steady accumulation of intracellular calcium around the MCV in U937 macrophages

**Supplementary video 3.** Homogeneous cytosolic distribution of intracellular calcium in U937 macrophages infected with Mm WT

**Supplementary video 4.** Intracellular calcium in U937 macrophages infected with Mm ΔRD1 mutant

**Supporting Table 1.** Reagents and Tools Table.

| Reagent/Resource | Reference or Source | Identifier or Catalog Number |
| --- | --- | --- |
| **Antibodies** |  |  |
| Anti-mono-and polyubiquitinated conjugates monoclonal antibody (FK2) (Monoclonal Mouse) | Enzo Life Science | Cat No. BML-PW8810-0100 |
| Anti-Galectin 3 (Rabbit Polyclonal) | Abcam | Cat No. ab53082 |
| Anti-Galectin 8 (Goat Polyclonal) | R&D Systems | Cat No. AF1305 |
| Anti-GAPDH (Polyclonal Rabbit) | Proteintech | Cat No. 10494-1-AP |
| Anti-GBP1 (Polyclonal Rabbit) | Proteintech | Cat No. 15303-1-AP |
| Anti-GBP1 (Monoclonal Mouse) | Proteintech | Cat No. 67161-1-Ig |
| Anti-GBP2 (Monoclonal Mouse) OTI5C8 | Novus | Cat No. NBP1-47768 |
| Anti-GBP4 (Polyclonal Rabbit) | Proteintech | Cat No. 17746-1-AP |
| Anti-LC3B (Polyclonal Rabbit) | MBL | Cat No. PM036 |
| Anti-LC3B (Polyclonal Mouse) | Cell Signalling | Cat No. 83506 |
| Anti-LAMP1 (Monoclonal Mouse) | Proteintech | Cat No. 67300-1-Ig |
| Anti-LAMP1 (Polyclonal Rabbit) | Abcam | Cat No. ab24170 |
| Anti-Cathepsin B (Polyclonal Rabbit) | Cell Signalling | Cat No. 31718 |
| Anti-TFEB (Polyclonal Rabbit) | Proteintech | Cat No. 13372-1-AP |
| **Chemicals, Enzymes and other reagents** |  |  |
| BAPTA-AM | Tocris | Cat No. 2787 |
| MLSA1 | Abcam | Cat No. ab144622 |
| Calmidazolium | TargetMol | Cat No. T10667 |
| Oregon Green 488 BAPTA-AM | MedChem | Cat No. HY-104058 |
| **Software** |  |  |
| GraphPad Prism | https://www.graphpad.com/scientific-software/prism/ | N/A |
| FlowJo | https://www.flowjo.com | N/A |
| Fiji | https://fiji.sc | N/A |
| **Other** |  |  |
| Cathepsin B Assay Kit (Magic Red) | Abcam | Cat No. ab270772 |
| LysoTracker™ Red DND-99 | Invitrogen™ | Cat No. L7528 |

## Supporting data 1. Uncropped blots with references bands indicating the area for each protein shown in the figures

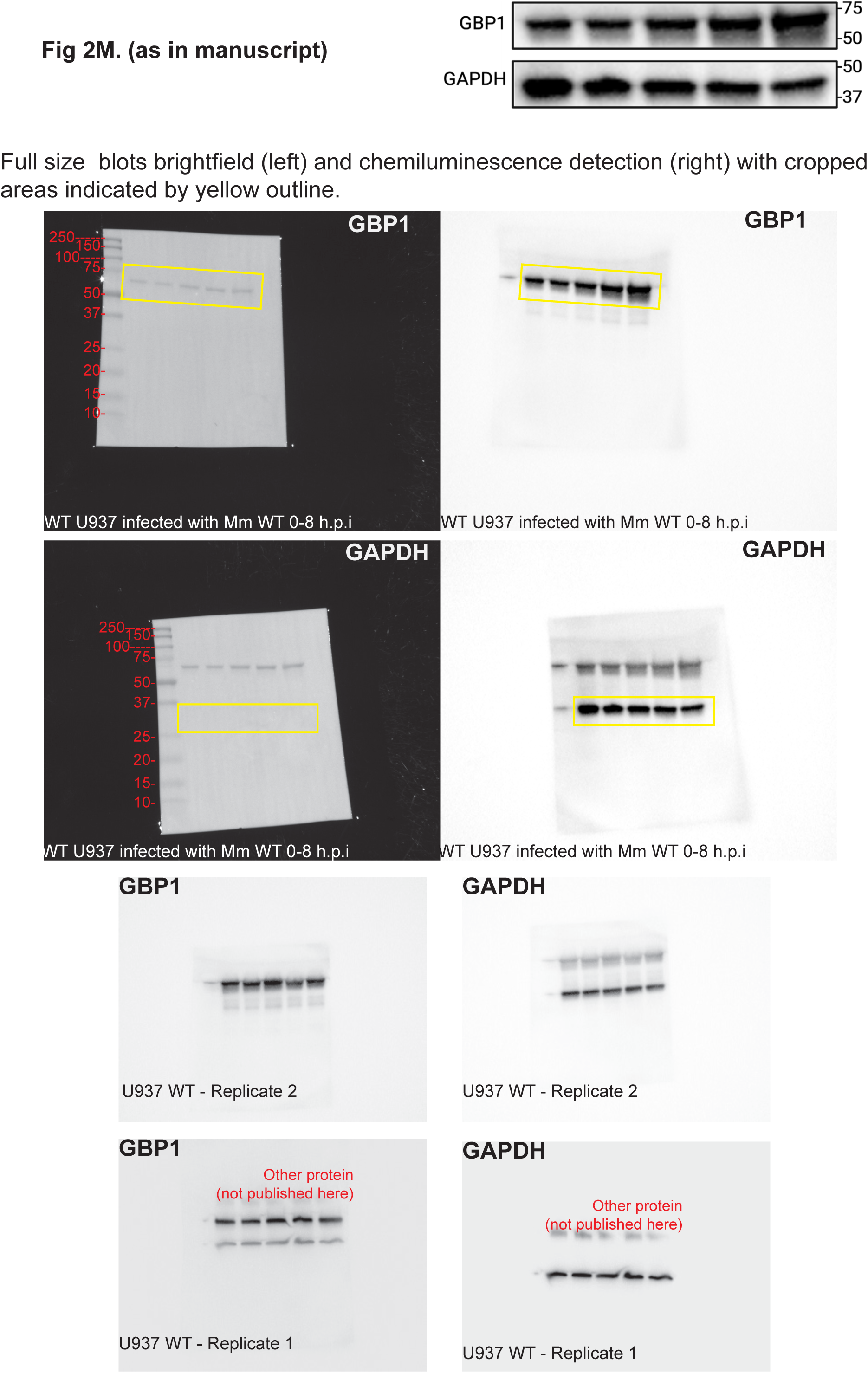
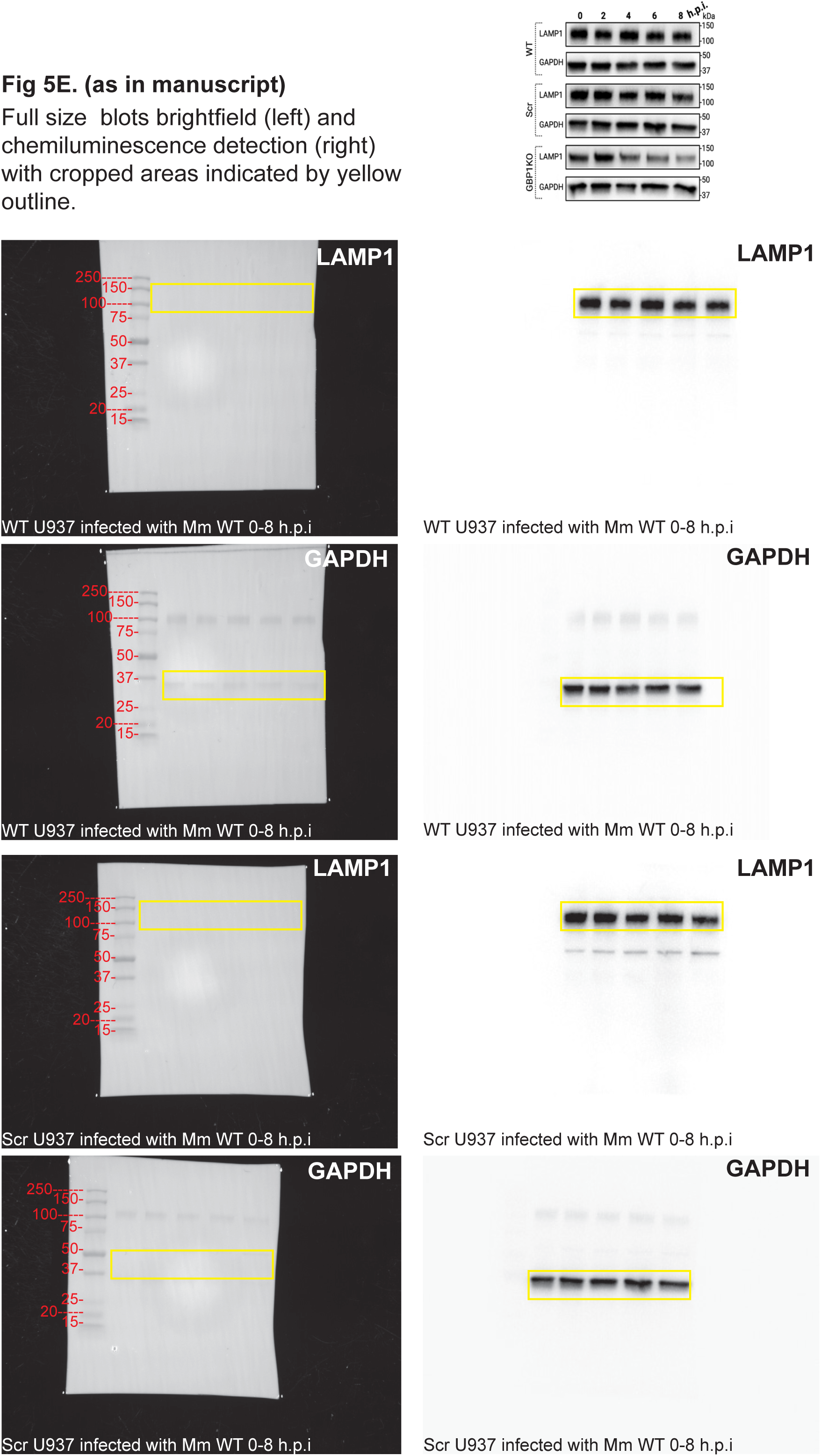
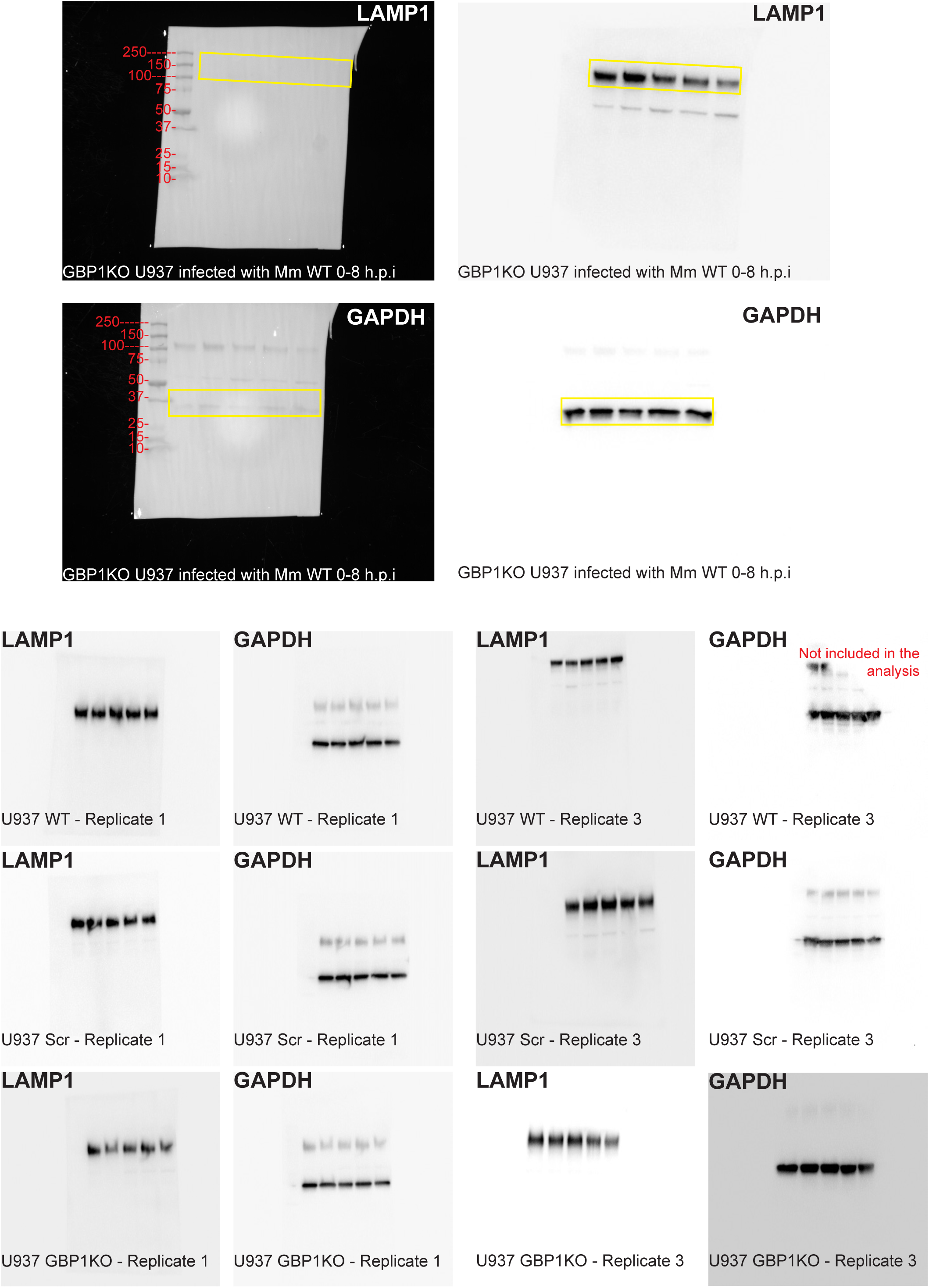
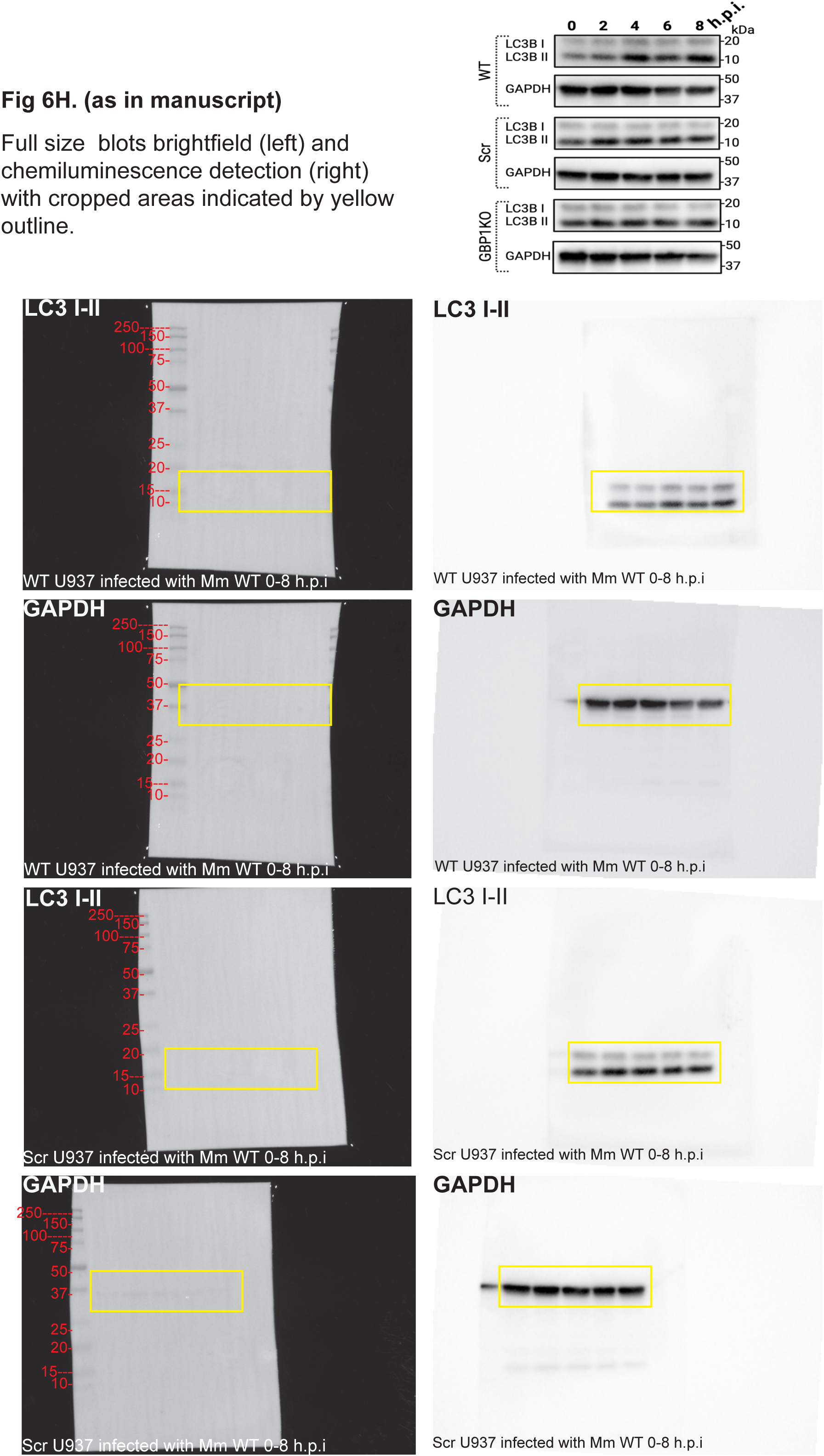
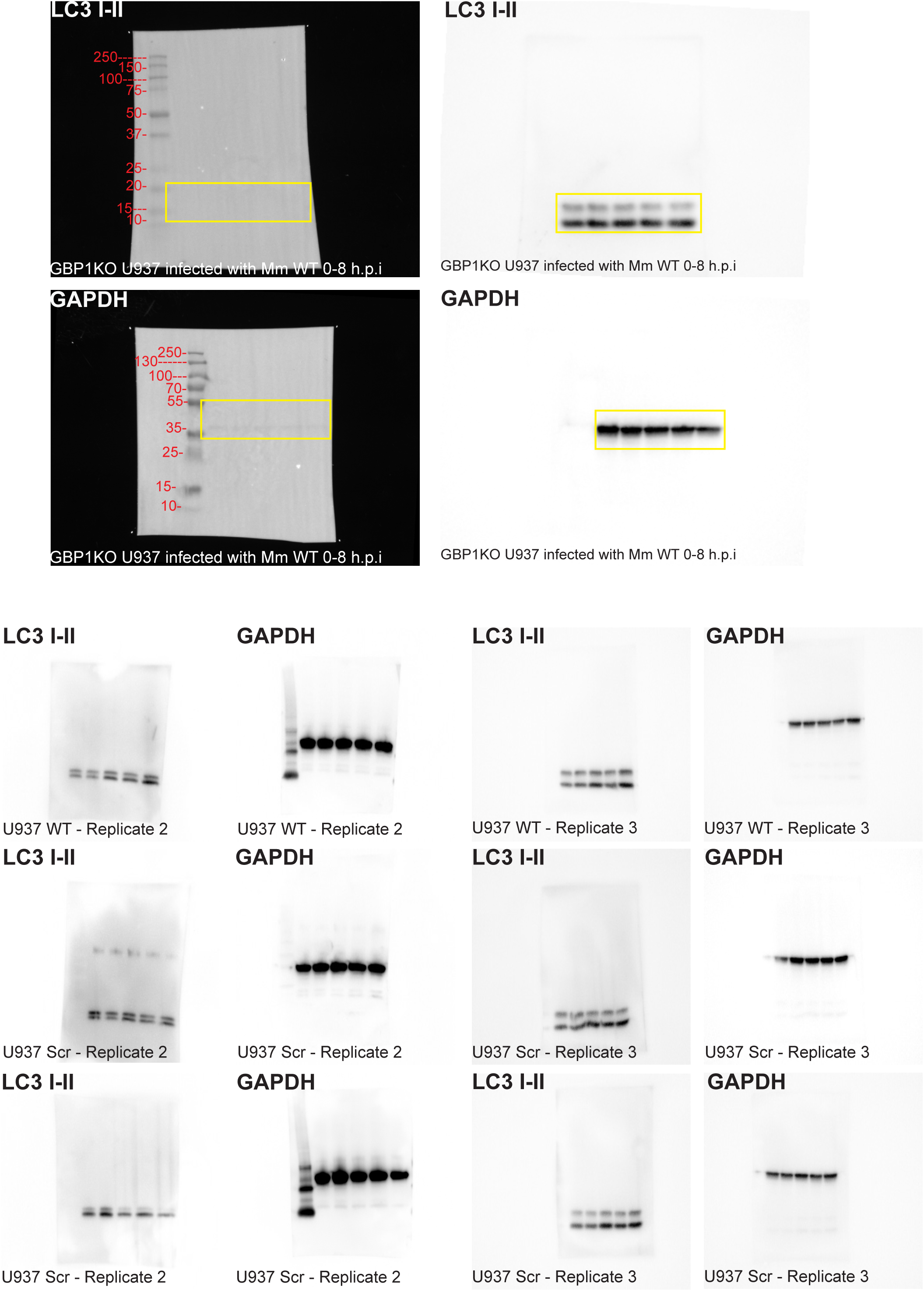
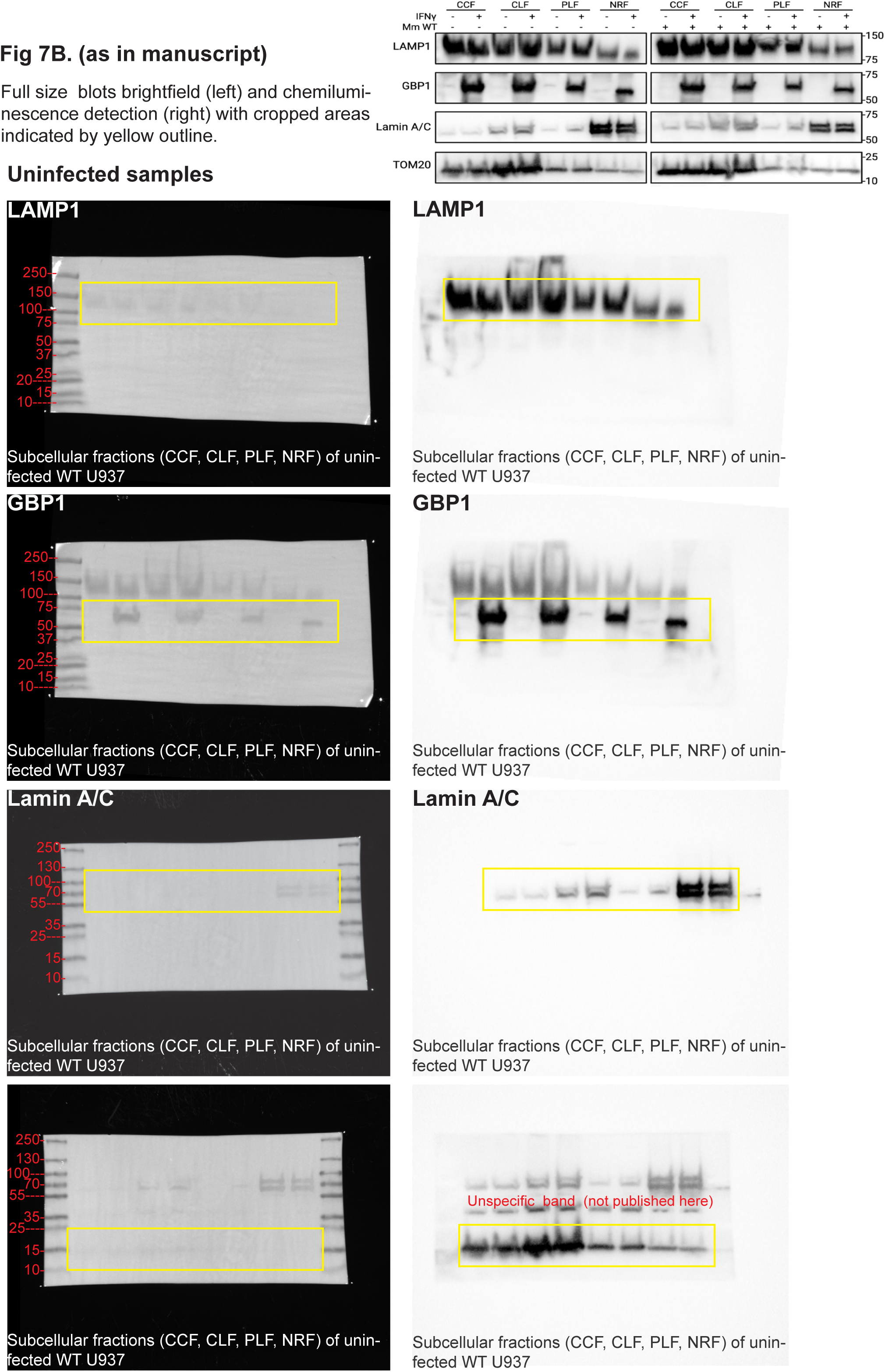
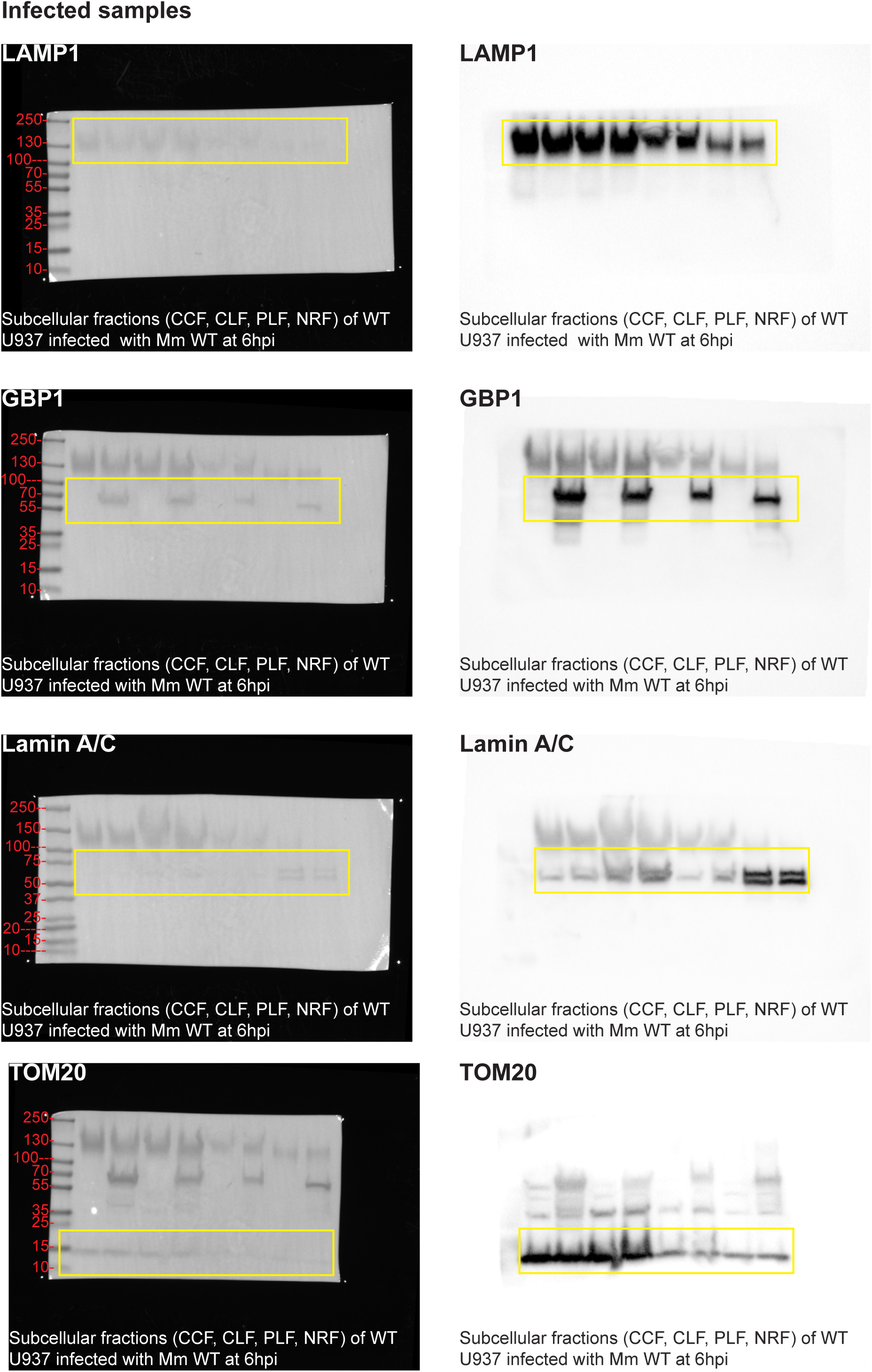
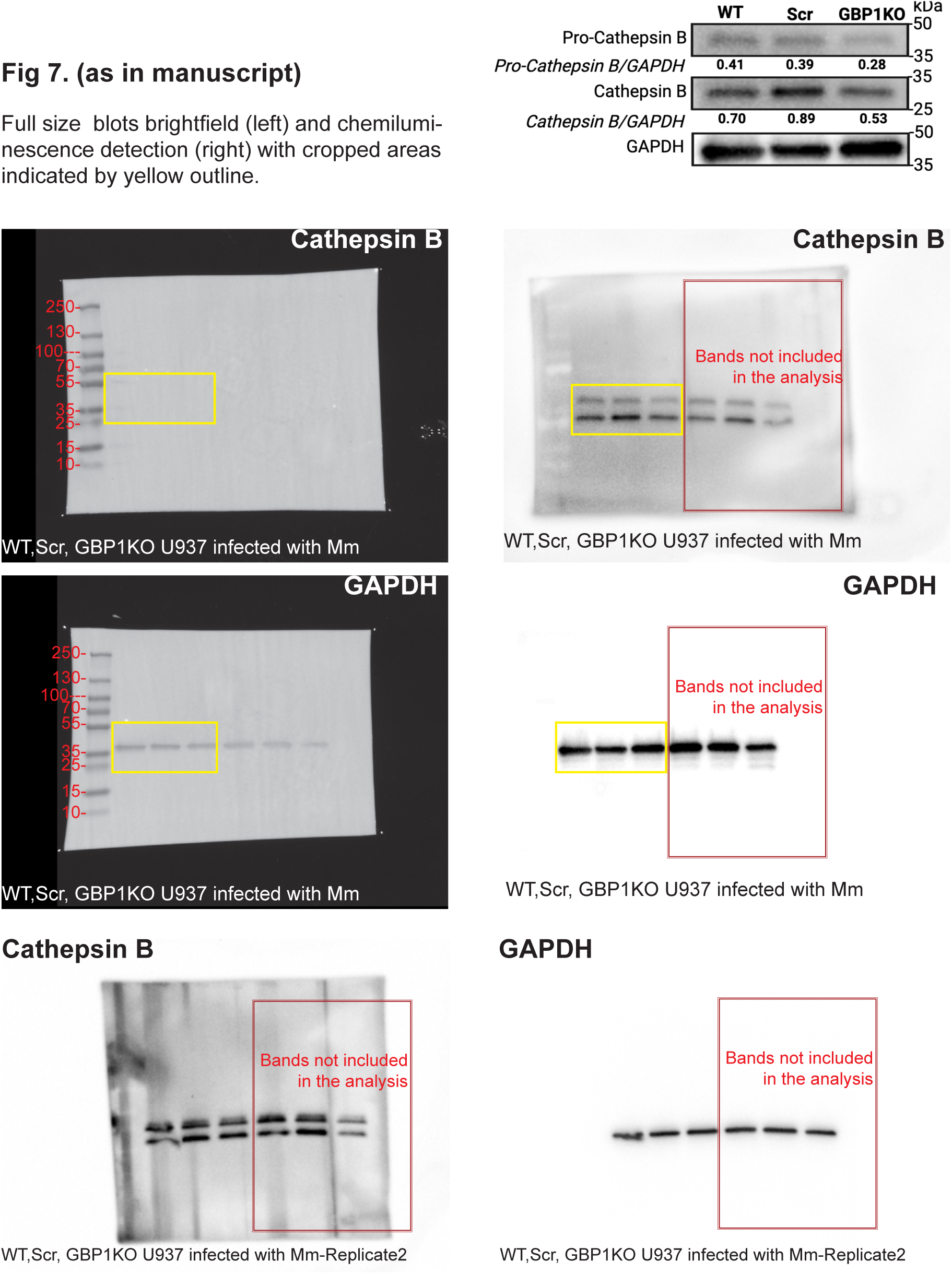

